# ICFinder: ion channel identification and ion permeation residue prediction using protein language models

**DOI:** 10.64898/2025.12.09.693328

**Authors:** Jue Wang, Xiaochun Zhang, Xiaoyu Fan, Bailong Xiao, Boxue Tian

**Affiliations:** MOE Key Laboratory of Bioinformatics, State Key Laboratory of Molecular Oncology, Beijing Frontier Research Center for Biological Structure, School of Pharmaceutical Sciences, Tsinghua University, Beijing, 100084, China

**Author notes:** To whom correspondence should be addressed: Boxue Tian.

**Keywords:** ion channel identification, ion permeation residue prediction, protein language models, webserver

## Abstract

Ion channel dysfunction underlies many diseases (e.g., arrhythmias, epilepsy, cystic fibrosis), and uncharacterized channels may also contribute to pathology. Identifying such channels and their residues directly contacting the permeation pathway (i.e., ion permeation residues) is key to elucidating transport mechanisms and developing targeted therapies. Leveraging the protein language model ESM-2 and curated datasets, we developed BLAPE and CLAPE frameworks for high-throughput ion channel identification and permeation residue prediction. Our models outperformed existing methods, with 33%-171% improvements in Matthews correlation coefficients (MCC) across different datasets. Analysis of amino acid composition revealed enrichment for weakly polar residues among ion permeation sites. Case studies on four diverse ion channels highlighted that CLAPE consistently outperforms existing predictors and remains applicable to proteins lacking experimental structures, while also complementing structure-based pipelines such as AlphaFold3. We further applied our models to UniRef50 to predict potential ion channels, and made these results publicly available through the ICFinder webserver (https://tianlab-tsinghua.cn/icfinder/), providing a ready-to-use resource for the research community. All source code is available at https://github.com/JueWangTHU/ICFinder.

## Introduction

Ions, such as K^+^, Na^+^, Ca^2+^, and Cl^−^, are essential components for a wide variety of physiological processes, including nerve impulse transmission, signal transduction, and muscle contraction, and therefore their transport into or out of cells and organelles is tightly regulated by ion channels (1–3). In 2004, Yu and colleagues constructed an evolutionary tree of ion channels based on sequence similarity (4), which included voltage-gated potassium and calcium channels, while numerous other ion channels were discovered in the following 20 years, such as the mechanosensitive PIEZO channels (5) or the TMEM63C monomeric ion channel (6). A typical channel consists of extracellular, transmembrane, and intracellular functional regions. For example, in calcium channels facilitating extracellular-to-intracellular ion permeation, calcium ions are initially attracted to residues in the extracellular region (7). Upon channel opening, ions traverse the transmembrane pore and exit near residues on the intracellular side.

Dysfunction or dysregulation of ion channels is closely associated with a variety of diseases, including cardiac arrhythmias such as long QT syndrome, neurological disorders such as epilepsy and pain disorders, and cystic fibrosis (8–10). Therefore, identifying previously uncharacterized ion channels and predicting their ion permeation residues can expand our understanding of disease-relevant ion transport mechanisms and provide a basis for the development of ion channel-targeted therapeutics. At present, electrophysiological techniques are the standard method for identifying ion channels, while pharmacological and immunological approaches are commonly employed to investigate their functions (11). In such pharmacological approaches, specific agonists, antagonists, or inhibitors are used to modulate ion channel activity and thereby determine their selectivity and sensitivity (12). Alternatively, immunological methods (e.g., immunohistochemistry, Western blotting, etc.) are commonly used to investigate ion channel expression, localization, and interactions with other molecules (13). Additionally, several innovative and well-established structural biology approaches, like X-ray crystallography, nuclear magnetic resonance (NMR), and cryo-electron microscopy (Cryo-EM), can be used to obtain high-resolution, atomic-level, three-dimensional (3D) structures of ion channels, facilitating investigation of their transport mechanisms and molecular interactions (14–16). For example, the classical structure of voltage-gated ion channels features S1-S6 domains, with S1-S4 acting as the voltage sensor, and S5-S6 forming the central pore responsible for ion permeation.

However, traditional experimental approaches are often labor-intensive, time-consuming, and costly, which makes the discovery of previously uncharacterized ion channels particularly challenging. In contrast, computational methods for identifying ion channels and predicting ion permeation residues are more convenient and faster. In particular, recent developments in deep learning and computational biology have further enhanced our ability to predict and analyze structure-function relationships of ion channels. The ion channel identification model, DeepPLM_mCNN (17) uses Evolutional Scale Modeling-1b (ESM-1b) (18) to extract sequence features, while and a multi-window Convolutional Neural Network (mCNN) (19) predicts ion channels from among candidate membrane proteins. Although no consensus method is yet available for predicting ion permeation residues, the TMP-MIBS model (20) can predict metal-binding sites in membrane proteins by combining position-specific scoring matrices (PSSM) (21) and physicochemical properties (PCP) (22), with a sliding window strategy to capture the influence of surrounding amino acids. Similarly, the CaBind_MCNN model (23) was developed to predict metal-binding sites in ion channels and ion transporters using 1,024-dimensional features generated by ProtTrans (24) and the same mCNN backbone network as DeepPLM_mCNN. Nevertheless, these computational approaches still demonstrate relatively limited accuracy. Furthermore, data imbalance (i.e., uneven distribution of positive and negative samples) continues to present a considerable challenge (25) for predicting ion channels and permeation residues. In TMP-MIBS, downsampling (26) is employed to address data imbalance, but can lead to inflated evaluation metrics, suggesting that it should be applied with caution (27).

To overcome these shortcomings and accelerate ion channel screening and mechanistic studies, in the present study, we leveraged the pre-trained protein language model (PLM), Evolutional Scale Modeling-2 (ESM-2) (28), to develop the Bidirectional Long Short-Term Memory (Bi-LSTM) (29) and Pre-trained Encoder (BLAPE) framework for ion channel identification, and the Contrastive Learning and Pre-trained Encoder (CLAPE) framework for ion permeation residue prediction, using the contrastive learning loss, triplet center loss (TCL) (30), to mitigate data imbalance. We constructed four datasets, including the Ion Channel Identification (ICIdentification) and Calcium Ion Channel Identification (CaICIdentification) datasets to train the BLAPE-ICIdentification and BLAPE-CaICIdentification models, respectively, for identifying ion channels; and the Ion Channel Permeation (ICPermeation) and Calcium Ion Channel Permeation (CaICPermeation) datasets to train the CLAPE-ICPermeation and CLAPE-CaICPermeation models for predicting ion permeation residues. All four models outperformed existing tools, showing higher prediction accuracy in both ion channel and ion permeation residue prediction tasks. We incorporated these models into a publicly accessible platform, Ion Channel Finder (ICFinder, https://tianlab-tsinghua.cn/icfinder/), which allows prediction of potential ion channels and their ion permeation residues, and provides structural information with key residues highlighted for various model organisms. This platform offers a ready-to-use resource for researchers, facilitating exploration of unreported ion channels and supporting the development of ion channel-targeted therapies. Its flexible design also allows rapid adaptation to additional datasets.

## Materials and methods

### HOLE program and definition of ion permeation residues

The HOLE program (31) uses a Monte Carlo simulated annealing procedure to analyze ion channel structures from PDB data, generating a new structure file with the pore geometry. In detail, raw PDB files were first cleaned by removing ions, small molecules, and lipids. Pore structures were then analyzed using HOLE, and the central 80% of each pore was extracted to compute the average radius and define the central axis. Residues within an additional 2 Å beyond the average radius were defined as ion permeation residues. HOLE can be downloaded, and further details can be found at https://www.holeprogram.org/.

### MMseqs2

MMseqs2 (32) is used for clustering protein sequences or retrieving similar sequences from databases, offering high speed and efficiency when handling large datasets, making it well-suited for this study. It is worth noting that different clustering tools may compute slightly different similarity scores for the same sequence pair, though the differences are generally small. As a result, using different tools such as UCLUST (33) or BLAST (34) may lead to minor variations in the final dataset, typically differing by a few clusters. Detailed usage instructions for MMseqs2 can be found at https://github.com/soedinglab/mmseqs2.

### Dataset splitting strategy

We divided the dataset into training, validation, and testing sets in an 8:1:1 ratio. Specifically, sequences were grouped in sets of ten, with the first sequence of each group assigned to the testing set to ensure a sufficient number of sequences for reliable performance evaluation. The second sequence was placed in the validation set, while the remaining sequences were assigned to the training set. This corresponds to the first fold of a 9-fold split, which serves as the default fold. The remaining folds were generated in the same manner.

### ESM-2

ESM-2 (28) is a PLM based on the Transformer architecture (35), designed for learning residue-level representations and predicting protein structures and functions. In this study, we used esm2_t33_650M_UR50D, a model trained on UniRef50 (36) with 650 million parameters across 33 layers. We extract 1,280-dimensional per-residue embeddings from the 33rd layer for downstream analysis. ESM-2 accepts the 20 standard amino acids as well as the unknown residue X, allowing us to retain sequences containing ambiguous residues without modification. While ESM-2 supports fine-tuning, we keep its parameters frozen in this study due to its substantially larger size (tens to hundreds of times the number of parameters in the subsequent Bi-LSTM and MLP backbone models). Prior studies, such as CLAPE-DB, have shown that fine-tuning ESM-2 has a negligible impact on model performance. For further details and implementation, ESM-2 is available at https://github.com/facebookresearch/esm.

### Backbone model of BLAPE: Bi-LSTM

In the BLAPE framework, the backbone model is Bi-LSTM (29). Bi-LSTM is a powerful sequence modeling architecture that captures long-range dependencies in both forward and backward directions. It is particularly well-suited for protein sequence analysis, where contextual relationships between amino acids are critical for understanding structure and function.

For a batch of sequence embeddings, Bi-LSTM follows these steps:

1. Padding to uniform length: The model first determines the longest sequence in the batch and pads all shorter sequences with zeros to match this length. The resulting input shape is:

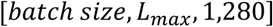

1. Bi-LSTM output: The model processes the sequences and outputs a hidden representation with the shape:

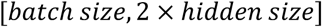

1. MLP classification: A simple MLP layer follows Bi-LSTM, transforming the hidden representation into a sequence-level probability prediction with the shape:

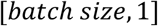

### Backbone model of CLAPE: MLP

In the CLAPE framework, the backbone model is a 5-layer MLP. The output dimensions of the five layers are 1,024, 256, 128, 64, and 2, respectively. Since each residue is treated as an independent sample, there is no need to handle variable-length sequences, making an MLP sufficient for this task.

### Triplet center loss

To enhance the separation between permeation and non-permeation residues, we adopt a contrastive learning loss named triplet center loss (30), which is expressed as:

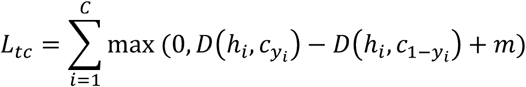

where 𝐷(ℎ_i_, 𝑐_yi_) and 𝐷(ℎ_i_, 𝑐_1-yi_) represent the Euclidean distances between the feature vector ℎ_i_ and the cluster centers of its own (𝑐_yi_) and opposite class (𝑐_1-yi_), respectively. Margin (𝑚) is a hyperparameter, which enforces a minimum separation between permeation and non-permeation residues.

Overall, in the CLAPE framework, the total loss function is formulated as:

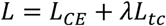

where lambda (𝜆), a hyperparameter, is referred to as the weight of the loss.

### Evaluation metrics

In this study, both the ground truth and predictions are represented as two equal-length, one-to-one corresponding binary sequences, where 0 denotes a negative sample and 1 denotes a positive sample. The four fundamental metrics used for evaluation are True Positive (TP), True Negative (TN), False Positive (FP), and False Negative (FN). TP represents cases where both the ground truth and prediction are 1, TN represents cases where both are 0, FP occurs when the ground truth is 0 but the prediction is 1, and FN occurs when the ground truth is 1 but the prediction is 0. Based on these, we primarily evaluate model performance using precision (Pre), recall (Rec), Matthews correlation coefficients (MCC), and area under the receiver operating characteristic curve (AUROC). Pre measures the proportion of correctly predicted positive samples among all predicted positives, while Rec quantifies the proportion of actual positive samples that are correctly identified. Precision and recall often trade off, and AUROC may be unreliable under data imbalance; therefore, we adopt MCC as the main evaluation metric (37,38). AUROC is used to assess the model’s overall ability to distinguish between positive and negative samples. The formulas for Pre, Rec, and MCC are as follows:

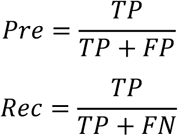

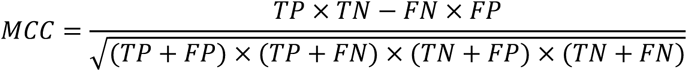

AUROC is computed using the scikit-learn Python package (https://scikit-learn.org/stable/).

### t-SNE dimension reduction

In this study, t-distributed Stochastic Neighbor Embedding (t-SNE) (39) was used to reduce the dimensionality of the 1,280-dimensional features embedded by ESM-2, the 512- and 2,048-dimensional features output by Bi-LSTM, and the 64-dimensional features generated by MLP, all projected into a 2D space to visualize the model’s ability to distinguish between positive and negative samples. The t-SNE implementation used in this study is from the scikit-learn package in Python, which can be downloaded and explored in more detail at https://scikit-learn.org/stable/.

### Integrated Gradients

To interpret how BLAPE distinguishes ion channels from other proteins, we adopted the Integrated Gradients (IG) method (40). IG attributes an importance score to each input feature by accumulating gradients of the model’s output along a straight-line path from a baseline input to the actual input. Formally, the attribution for input 𝑥_i_ is given by:

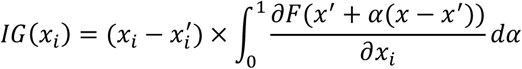

where 𝑥 is the embedding of the target protein sequence, 𝑥^i^ is the baseline, which is often set to an all-zero vector, and 𝐹 is the model.

In our study, we observed that Bi-LSTM architectures yield higher IG scores at sequence termini than in central regions. To correct for this positional bias, we generated *N* random protein sequences with the same length as the target sequence and computed their IG scores. The absolute values of these scores were averaged to obtain a positional background contribution. The normalized attribution for each residue was then calculated as the raw IG score divided by the background score at the same position. Then, residues with the highest normalized scores (top *k*) were considered potentially important for ion channel function. A larger *N* reduces estimation error but increases computational cost, whereas a larger *k* broadens the coverage of functional residues but also introduces less relevant positions. We set *N* = 5, *k* = 15% in the case study.

IG can be implemented in Python using the Captum library (https://captum.ai/).

### PyMOL visiualization

PyMOL is a powerful molecular visualization tool widely used in structural biology for displaying and analyzing 3D structures of biomolecules. In this study, PyMOL is employed to visualize ion channel structures. After selecting a UniProt ID, the corresponding PDB ID is identified, and the 3D structure is obtained using PyMOL’s “Get PDB” function. The “Display-Sequence” feature is then used to display the amino acid sequence. Depending on the analysis requirements, different colors and shapes are applied to highlight key regions, such as transmembrane segments and real or predicted ion permeation residues. For more information and to download PyMOL, visit https://pymol.org/.

### DeepTMHMM

DeepTMHMM (41) is an advanced deep-learning-based tool for predicting the topology of transmembrane proteins with high accuracy. It provides residue-level annotations, distinguishing signal peptides (S), transmembrane regions (M), extracellular regions (O), and intracellular regions (I). In this study, DeepTMHMM is used to predict residue-level topology information. For organelle membrane proteins, such as mitochondrial inner membrane proteins, O represents the intermembrane space, while I corresponds to the matrix. To use DeepTMHMM, visit https://services.healthtech.dtu.dk/services/DeepTMHMM-1.0/.

### Seq2Symm

Seq2Symm (42) is a deep learning-based tool designed to predict the symmetry type and oligomeric state of protein complexes directly from amino acid sequences. It classifies symmetry into categories such as cyclic (Cn), dihedral (Dn), and asymmetric (C1), and can also output uncertain predictions as “CX”, indicating ambiguity in the oligomeric state and often suggesting a higher-order assembly beyond a hexamer. Seq2Symm enables researchers to estimate plausible assembly states prior to structure prediction by tools like AlphaFold3, which require predefined symmetry and oligomerization as input. More details about Seq2Symm are available at https://github.com/microsoft/seq2symm.

### AlphaFold3

AlphaFold3 (43) is the latest version of DeepMind’s protein structure prediction model. In addition to high-accuracy predictions of monomeric and multimeric protein structures, AlphaFold3 introduces the ability to model protein-ligand, protein-DNA/RNA, and protein-small molecule interactions. It accepts not only amino acid sequences but also additional input such as oligomeric states, symmetry, and binding partners, making it particularly powerful for complex systems like membrane proteins and ion channels in our study. AlphaFold3 is available at https://alphafoldserver.com/.

### Computational resources and efficiency

All experiments were conducted on two GPU setups: four NVIDIA A100-PCIE-40GB GPUs with an Intel Xeon Silver 4210R CPU @ 2.40 GHz, and eight NVIDIA Tesla V100-SXM2-32GB GPUs with an Intel Xeon Gold 6226R CPU @ 2.90 GHz. Multi-GPU parallelization was employed to accelerate 9-fold cross-validation and hyperparameter tuning. ESM-2 embeddings for sequences shorter than 1,024 residues were generated on GPUs, allowing tens of sequences per second, while longer sequences were computed on CPUs due to GPU memory limitations, with a single sequence taking approximately 10 seconds; such long sequences were rare. Model training was performed entirely on GPUs. CLAPE, which incorporates TCL, required slightly less than 1 second per sequence per epoch, whereas BLAPE, using cross-entropy (CE) loss, processed hundreds of sequences per second. Inference and testing were conducted on CPUs, where hundreds of sequences could be processed in less than a second. Computational efficiency for different tasks is summarized in Supplementary Table 1.

## Results

### Curation of Ion channel identification and ion permeation residue prediction datasets

To train a model for ion channel identification in public sequencing databases, we constructed the ICIdentification dataset based on the previously published ExpGO65 dataset (44). The original ExpGO65 dataset contained 65 binary labels per protein, indicating the presence or absence of 65 common Gene Ontology (GO) terms (45), including “channel activity” (GO:0015267). Since our study focused on ion channels, only proteins with “monoatomic ion channel activity” GO terms (GO:0005216) were assigned a positive label, while all other sequences were assigned a negative label (Figure 1A). The ICIdentification dataset was then clustered on 50% sequence similarity threshold using MMseqs2 (32), yielding 33,765 clusters. After selecting the cluster center of each cluster as representative sequences, 80% of the representative sequences were randomly partitioned into training, validation, and testing sets in an 8:1:1 ratio. The remaining 20% of representative sequences were reserved as the independent testing set. To investigate whether he ion selectivity of ion channels can be predicted, we curated the CaICIdentification dataset to train a model capable of distinguishing calcium channels from other proteins. This dataset included amino acid sequences of 29 calcium ion channels, 43 non-calcium channels, and additional non-ion channel proteins obtained from the ICIdentification dataset. The same clustering and partitioning strategy were applied to generate training, validation, and testing sets.

**Figure 1.**
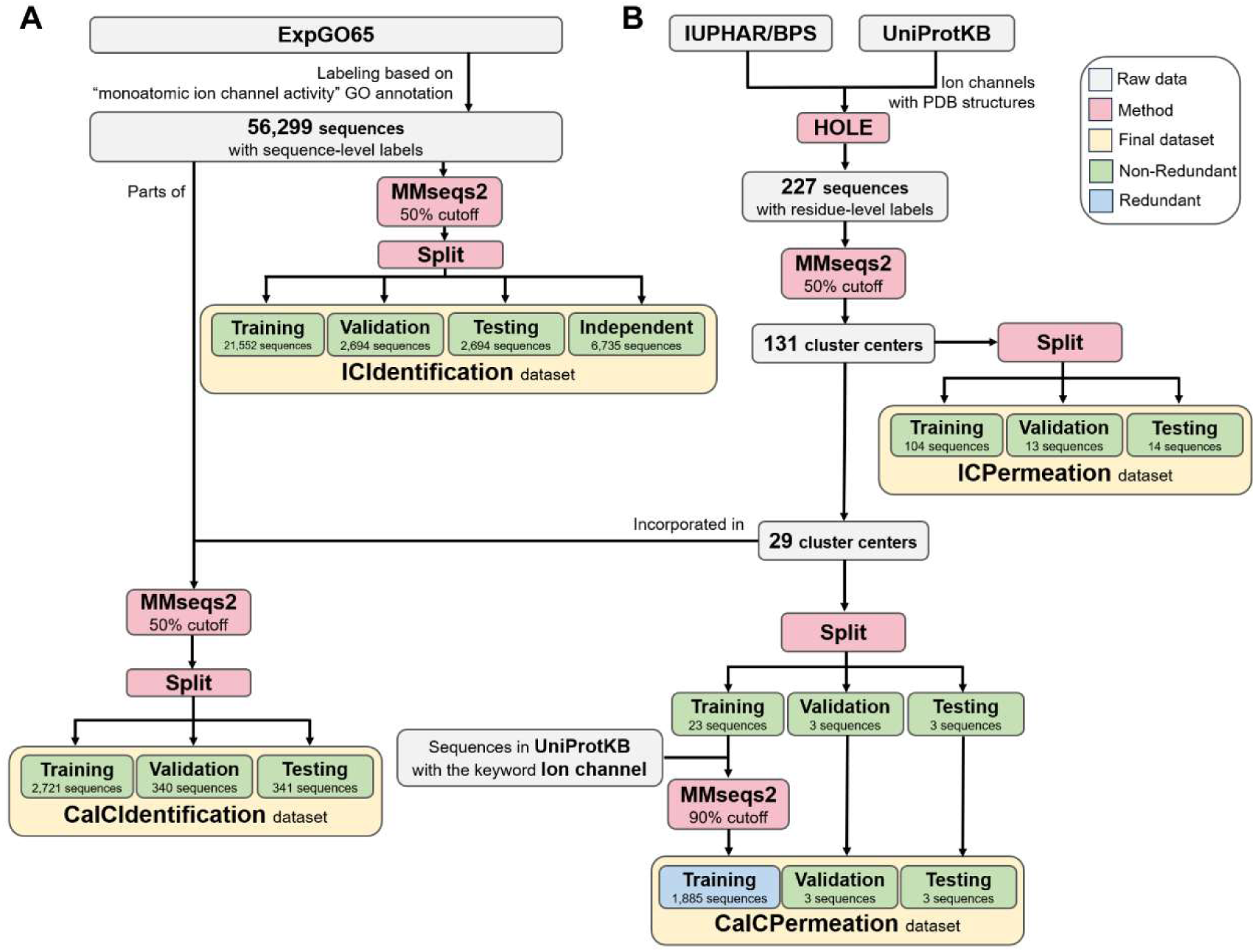
Dataset curation for constructing ion channel identification (**A**) and ion permeation residue prediction (**B**) models. (**A**) All protein sequences were relabeled based on the presence or absence of the GO term “monoatomic ion channel activity”, resulting in the ICIdentification dataset. This dataset was subsequently split into training, validation, and testing sets, along with an independent testing set. To construct CaICIdentification, we balanced positive samples (29 calcium channels), ambiguous 43 non-calcium channels, and negative samples (3,330 non-ion channels). (**B**) PDB sequences annotated to contain ion channel structures were collected for analysis with HOLE program to identify ion permeation residues. In total, 227 such sequences were collected. Clustering them using MMseqs2 with a 50% identity cutoff generated 131 cluster centers, which were then split into non-redundant training, validation, and testing sets. A subset of 29 calcium channels was selected from the 131 cluster centers and processed similarly. To assemble the CalCPermeation dataset, comprising only calcium channels, the training set was expanded with ≥90% similarity sequences to form a redundant set of 1,885 sequences, while the validation and testing sets remained non-redundant.

Additionally, to construct a model capable of identifying ion permeation residues, we curated the ICPermeation and CaICPermeation datasets (Figure 1B). The process for assembling ICPermeation followed two stages, in which ion channels with PDB structures were first collected from the IUPHAR/BPS Guide to Pharmacology (46) and UniProt databases (47). We then applied HOLE program (31) to identify channel pores. Residues located within a distance from the pore center equal to the average pore radius plus 2 Å were labeled as ion permeation residues; a binary label was also assigned to the full sequence, where “1” indicated permeation residues and “0” denoted non-permeation residues. All identified residues were manually inspected, and errors in structure identification were corrected. This process yielded 227 total sequences. To further expand the dataset, these sequences were clustered using MMseqs2 with a 50% similarity threshold, which resulted in a set of 131 clusters. The cluster centers were partitioned into non-redundant training, validation, and testing sets at an 8:1:1 ratio.

To specifically identify calcium ion permeation residues, we constructed the CaICPermeation subset. Examination of UniProt descriptions and GO annotations of the 131 ICPermeation clusters uncovered 29 experimentally validated or putative calcium ion channels that could serve as the basis for dataset construction following the same process as for ICPermeation. However, as the training set contained only 23 sequences, which is inadequate to build an accurate model, we added sequences from UniProt that shared at least 90% similarity to the training set, thus expanding CaICPermeation to 1,885 sequences. Detailed information for these four datasets (e.g., positive and negative sample numbers, average sequence length) is provided in Table 1.

**Table 1.** Summary of four datasets used in this study

|  |  | Num | Pos | Neg | % of Pos | Av. Length (AA) |
| --- | --- | --- | --- | --- | --- | --- |
| ICIdentification | Training | 21,552 | 284 | 21,268 | 1.32 | 424 |
|  | Validation | 2,694 | 42 | 2,652 | 1.56 | 420 |
|  | Testing | 2,694 | 38 | 2,656 | 1.41 | 418 |
|  | Independent | 6,735 | 97 | 6,638 | 1.44 | 422 |
| CaICIdentification | Training | 2,721 | 22 | 2,699 | 0.81 | 431 |
|  | Validation | 340 | 2 | 338 | 0.59 | 421 |
|  | Testing | 341 | 5 | 336 | 1.47 | 445 |
| ICPermeation | Training | 104 | 2,458 | 71,213 | 3.34 | 708 |
|  | Validation | 13 | 263 | 9,701 | 2.64 | 766 |
|  | Testing | 14 | 300 | 12,121 | 2.42 | 887 |
| CaICPermeation | Training | 1,885 | 37,573 | 1,593,057 | 2.30 | 865 |
|  | Validation | 3 | 51 | 1,566 | 3.15 | 539 |
|  | Testing | 3 | 63 | 3,165 | 1.95 | 1,076 |

### BLAPE and CLAPE model architecture

We developed the BLAPE (Figure 2A) and CLAPE (Figure 2B) frameworks to enable binary classification of ion channel proteins and ion permeation residues, respectively, relying solely on sequence features. In both models, input protein sequences were first transformed into 1,280-dimensional feature vectors for each residue using the ESM-2. To prevent excessive resource consumption and mitigate risk of catastrophic forgetting (48,49), the ESM-2 parameters were not changed during model training throughout training. BLAPE was then used to process the matrix of the full sequence length (L) × 1,280 features using a Bi-LSTM network, with padding applied to handle sequences of varying lengths. The intermediate feature representations provided by Bi-LSTM were subsequently fed into a simple 1-layer MLP with a Sigmoid activation function to generate sequence-level classifications. BLAPE was trained on the ICIdentification and CaICIdentification datasets to establish the BLAPE-ICIdentification and the BLAPE-CaICIdentification models, respectively.

**Figure 2.**
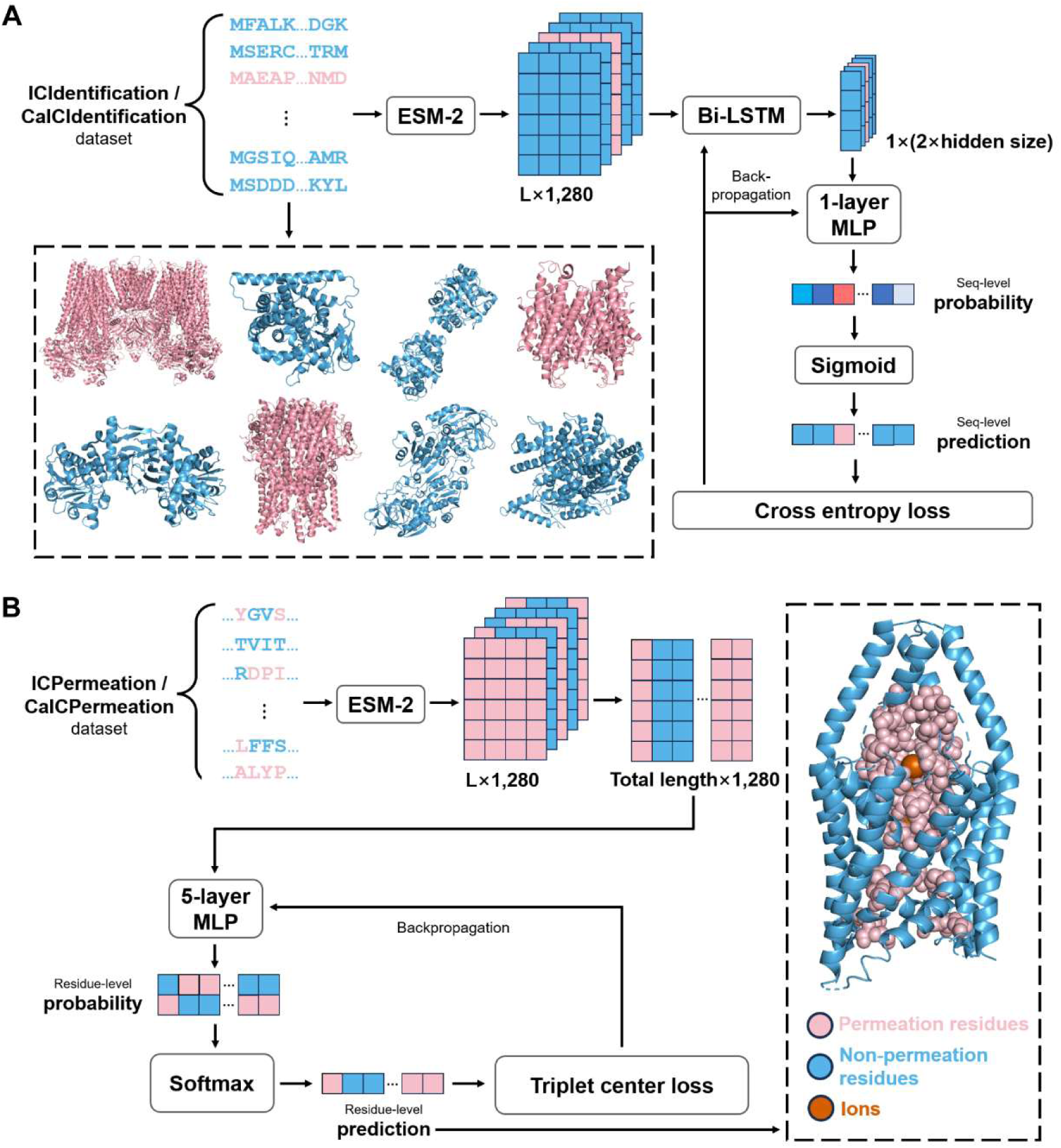
Overall model architecture of BLAPE and CLAPE. (**A**) BLAPE requires a sequence-level labeled dataset (pink, ion channels; blue, non-ion channels). Each sequence is processed by ESM-2, generating L × 1,280 features (L varies across sequences). Extracted features are passed through a Bi-LSTM, then a single-layer MLP with a Sigmoid activation function to produce sequence-level predictions. BLAPE is only trained with CE loss, and only the Bi-LSTM and MLP modules are trainable. (**B**) CLAPE requires a residual-level labeled dataset (pink, permeation residues; blue, non-permeation residues). All sequences are processed by ESM-2, and the resulting features are concatenated into a large-scale matrix that has a size of total sequence length × 1,280 features, with. each column of the matrix representing an individual sample. A trainable 5-layer MLP with a Softmax activation function finally generates residual-level predictions. TCL is incorporated to improve model performance.

Alternatively, CLAPE was built to view the L × 1,280 feature matrix as L individual samples and classify each residue separately using a five-layer MLP, with the final Softmax layer producing residue-level predictions. Training CLAPE on the ICPermeation and CaICPermeation datasets resulted in the CLAPE-ICPermeation model for general prediction of ion permeation residues, while training on the CaICPermeation dataset generated the CLAPE-CaICPermeation model for specifically predicting calcium ion permeation residues. Examination of loss functions showed that TCL only improved model performance within the CLAPE framework, although both datasets had imbalanced data. Based on these results, BLAPE was trained using only simple CE loss.

### Performance evaluation with existing models

To evaluate the performance of our models, we also trained previously reported models for ion channel identification and permeation residue prediction using our datasets to allow direct comparison. All experiments were conducted using five, independent, random seeds, and the final evaluation metrics were obtained by averaging performance across these seeds (Supplementary Table 2). Comparing BLAPE-ICIdentification with DeepPLM_mCNN in an ion channel binary classification task, we found that our model achieved an MCC of 0.740 on the ICIdentification dataset, outperforming DeepPLM_mCNN (MCC = 0.415, Table 2). BLAPE-ICIdentification also reached an MCC of 0.626 and an AUROC of 0.935 on the independent testing set (Supplementary Table 3). Similarly, BLAPE-CaICIdentification attained an MCC of 0.936 on the CaICIdentification dataset, representing a 171% relative improvement over DeepPLM_mCNN (MCC = 0.346, Table 3). These results indicated that BLAPE could identify ion channels with high accuracy.

**Table 2.** Comparison of BLAPE-ICIdentification with DeepPLM_mCNN on the ICIdentification dataset

|  | Pre | Rec | MCC | AUROC |
| --- | --- | --- | --- | --- |
| BLAPE-ICIdentification | 0.920±0.035 | 0.600±0.031 | **0.740±0.026** | 0.962±0.005 |
| DeepPLM_mCNN | 0.194±0.042 | 0.963±0.021 | 0.415±0.042 | **0.987±0.001** |

**Table 3.** Comparison of BLAPE-CaICIdentification with DeepPLM_mCNN on the CaICIdentification dataset

|  | Pre | Rec | MCC | AUROC |
| --- | --- | --- | --- | --- |
| BLAPE-CaICIdentification | 1.000±0.000 | 0.880±0.098 | **0.936±0.052** | **1.000±0.000** |
| DeepPLM_mCNN | 0.182±0.155 | 0.933±0.094 | 0.346±0.152 | 0.947±0.015 |

As we could find no previous studies directly addressing the prediction of ion permeation residues, we benchmarked our CLAPE-ICPermeation model against CaBind_MCNN, which targets calcium ion-binding sites in calcium channels, and TMP-MIBS, which predicts ion-binding sites in membrane proteins. As these models employ multi-window scanning strategies, their features are often redundantly encoded, leading to considerably higher memory consumption compared to our approach. As the ICPermeation dataset comprised only 131 sequences, we followed the original GitHub implementation and applied the full multi-window configuration. Testing on the ICPermeation dataset, our CLAPE-ICPermeation model achieved an MCC of 0.449, surpassing both CaBind_MCNN (MCC = 0.220) and TMP-MIBS (MCC = 0.283), while its AUROC (0.842) was between the two (0.871 and 0.556, respectively). Closer examination of precision and recall revealed that CaBind_MCNN exhibited high recall but low precision, while TMP-MIBS demonstrated high precision but low recall (Table 4). Comparison of predictions on the CaICPermeation dataset showed similar results, with CLAPE-CaICPermeation achieving the highest MCC (0.694) (Table 5). Consistent results were obtained in 9-fold cross-validation across all four datasets (Supplementary Table 4), with small standard deviations indicating stable model performance. Collectively, these analyses showed that the CLAPE framework could outperform other existing models across the evaluated datasets.

**Table 4.** Comparison of CLAPE-ICPermeation with other models on the ICPermeation dataset

|  | Pre | Rec | MCC | AUROC |
| --- | --- | --- | --- | --- |
| CLAPE-ICPermeation | 0.699±0.053 | 0.301±0.044 | **0.449±0.028** | 0.842±0.011 |
| CaBind-MCNN | 0.089±0.013 | 0.798±0.023 | 0.220±0.020 | **0.871±0.003** |
| TMP-MIBS | 0.743±0.051 | 0.112±0.006 | 0.283±0.018 | 0.556±0.003 |

**Table 5.** Comparison of CLAPE-CaICPermeation with other models on the CaICPermeation dataset

|  | Pre | Rec | MCC | AUROC |
| --- | --- | --- | --- | --- |
| CLAPE-CaICPermeation | 0.751±0.016 | 0.651±0.018 | **0.694±0.010** | **0.976±0.003** |
| CaBind-MCNN | 0.232±0.092 | 0.852±0.033 | 0.416±0.080 | 0.961±0.005 |
| TMP-MIBS | 0.821±0.014 | 0.339±0.007 | 0.522±0.006 | 0.669±0.003 |

To examine the ability of BLAPE-ICIdentification, BLAPE-CaICIdentification, CLAPE-ICPermeation, and CLAPE-CaICPermeation to distinguish positive and negative samples, we applied t-SNE (39) dimensionality reduction on the respective datasets (Figure 3). For BLAPE models based on Bi-LSTM, we extracted the hidden representations from the model output, while using intermediate features from the fourth layer for MLP-based CLAPE models. For comparison, we also applied t-SNE to the raw ESM-2-generated features. The results indicated that our trained models could effectively cluster positive samples together, whereas the raw features contained interspersed positive and negative samples.

**Figure 3.**
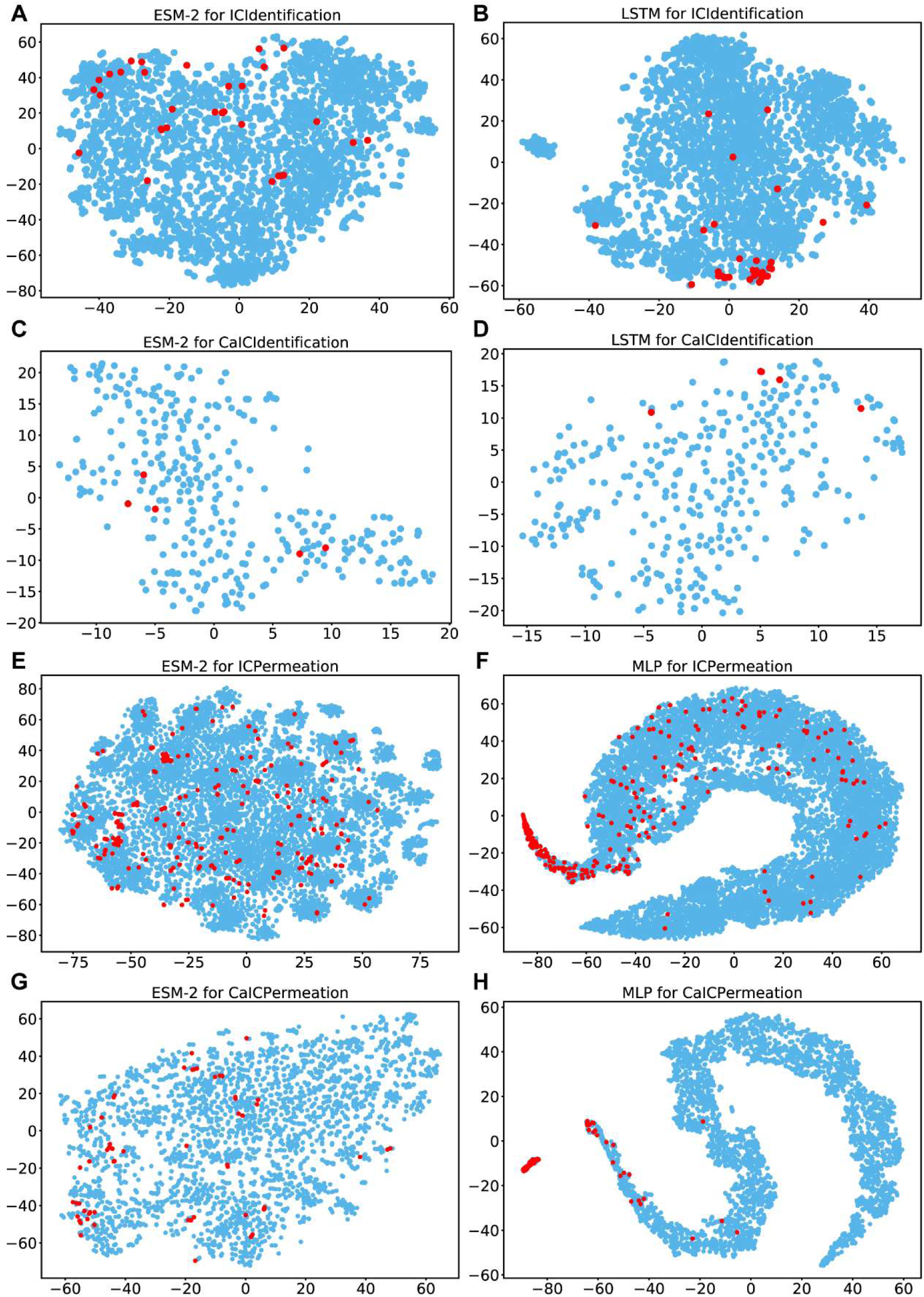
Dimensionality reduction of raw and learned representations. (**A, B**) t-SNE plots comparing of the ICIdentification dataset, while (**C, D**) correspond to the CaICIdentification dataset. (**E, F**) depict the ICPermeation dataset, and (**G, H**) illustrate the CaICPermeation dataset. In all plots, red dots indicate positive samples, and blue dots represent negative samples. (**A, C**) show the t-SNE projection of hidden representations obtained from Bi-LSTM, while (**E, G**) visualize the fourth-layer 64-dimensional output from the MLP. The well-trained models effectively separate positive and negative samples. In contrast, (**B, D, F, H**) display the t-SNE results of the original 1280-dimensional ESM-2 embeddings, where positive and negative samples remain largely intermixed.

### Hyperparameter optimization to improve model performance

To improve the prediction accuracy of our models, we sought to optimize the hyperparameters by first determining the most effective strategy for embedding protein sequences into features. Evaluating the performance of two widely used PLMs, ESM-2 (28) and ProtBERT (24), across all four datasets under the BLAPE and CLAPE frameworks (Supplementary Table 5) demonstrated that ESM-2 consistently outperformed ProtBERT, with approximately 27% higher average MCC and a roughly 2% increase in AUROC. The superior performance of ESM-2 was likely due to its considerably larger number of parameters (650 million vs. 110 million) and the higher dimensionality of its feature representations (1,280 dimensions vs. 1,024 dimensions).

Beyond the high-dimensional embeddings from PLMs, we conducted comparison of performance on traditional amino acid features, such as Atchley factors (50) and protein-protein interaction (PPI) probabilities, but the results were not promising. Derived and simplified from the ∼500-dimensional AAindex (22), Atchley factors are 5-dimensional amino acid features that describe the five main properties of amino acids, including polarity, secondary structure, molecular size, codon frequency, and electrostatic charge. As these features do not capture evolutionary information or positional context within the sequence, we concatenated Atchley factors with ESM-2 embeddings, which resulted in 1,285-dimensional features. However, their inclusion had minor effect on both MCC and AUROC on the ICPermeation and CaICPermeation datasets (Supplementary Figure 1). These findings suggested that ESM-2 embeddings may implicitly encode fundamental amino acid properties, making Atchley factors redundant.

Given that most ion channels function as multimers and typically contain at least two homologous subunits, we hypothesized that training with PPI information might facilitate prediction of ion permeation residues, as these residues tend to be located at interfaces rather than deeper, core regions. To incorporate PPI information, we predicted residue-level PPI scores, which are scalar values assigned to each residue. These scores were added to every dimension of the ESM-2 embeddings, generating a fused feature. We then trained a model on a homologous PPI dataset (44) to predict PPI sites of ion channels. Incorporating PPI predictions improved MCC on the ICPermeation but reduced it on the CaICPermeation, likely due to the modest accuracy of our PPI model (MCC = 0.388) and task-specific differences. These results indicate that PPI features may be useful, but their utility depends on both better predictor quality and downstream context.

Next, we evaluated the contribution of the contrastive learning loss, namely TCL (30) in our study, to the CLAPE framework. On the ICPermeation dataset, we compared model performance with or without TCL, using either CE loss or focal loss (51), another method to mitigate data imbalance (Table 6). The results showed that CE loss alone without TCL achieved the lowest MCC (0.418). Incorporating focal loss or TCL individually increased MCC by 5.7% and 7.4%, respectively, reaching 0.442 and 0.449. However, combining both losses led to a reduction in MCC to 0.432. This decrease likely reflects partially redundant emphasis on negative samples during training, which may limit the model’s generalization performance on the testing set. Based on these observations, we selected TCL with standard CE loss for training the CLAPE models, balancing improved performance with robust generalization.

**Table 6.** Model performance with different loss functions on the ICPermeation dataset

| Loss function |  | MCC | AUROC |
| --- | --- | --- | --- |
| CE loss | -TCL | 0.418±0.010 | 0.855±0.006 |
|  | +TCL | **0.449±0.028** | 0.842±0.011 |
| Focal loss | -TCL | 0.442±0.019 | **0.861±0.007** |
|  | +TCL | 0.432±0.021 | 0.848±0.004 |

The third stage of optimization focused on feature transformation for final predictions. We conducted framework-specific hyperparameter tuning by optimizing Bi-LSTM architecture for BLAPE and adjusting loss function for CLAPE. For BLAPE-ICIdentification, hyperparameter tuning (Figure 4A-D) revealed that the strongest configuration included a single LSTM layer with a hidden size of 256 and a dropout of 0.3. Additionally, we tuned the learning rate, ultimately choosing 5 × 10⁻^5^. Using the same method, the BLAPE-CaICIdentification model performed better with a hidden size of 1,024 and a learning rate of 1 × 10⁻^4^. For CLAPE-ICPermeation (Figure 4E-H), the optimal hyperparameters were identified as margin = 5, λ = 1.0, with learning rates set to 0.01 for TCL and 1 × 10⁻⁴ for the classifier. Similarly, we found the strongest hyperparameters for CLAPE-CaICPermeation were margin = 12, λ = 0.1, and the same learning rates. Finally, comparison of performance between MLP and CNN architectures in the CLAPE framework (Supplementary Figure 2) showed that both architectures were viable. Notably, both CNN (CLAPE-DB (52)) and MLP (CLAPE-SMB (53)) backbones have been effective in prior CLAPE-based tasks. **Table 6**. Model performance with different loss functions on the ICPermeation dataset CE loss Focal loss

**Figure 4.**
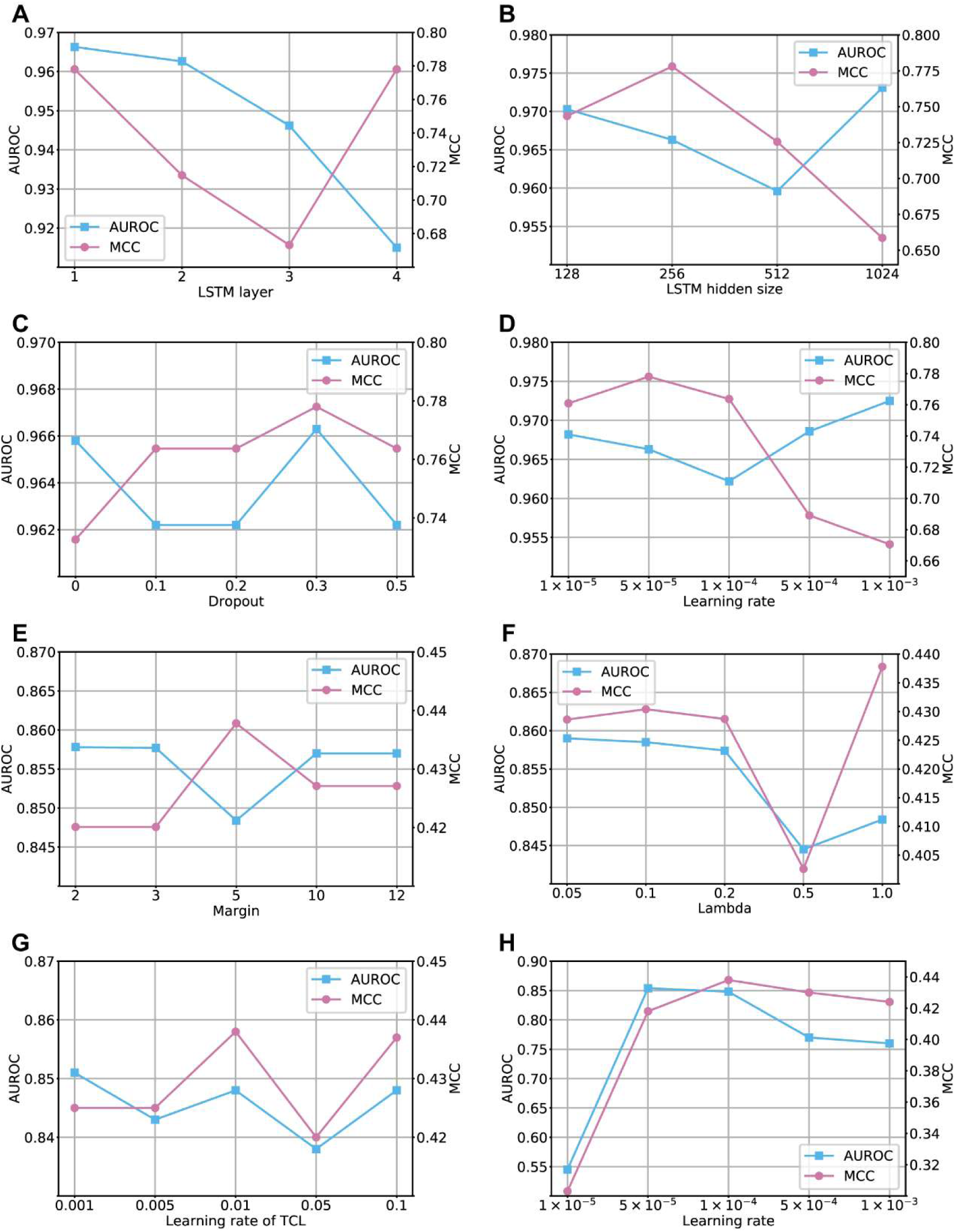
Hyperparameter optimization for BLAPE-ICIdentification (**A-D**) and CLAPE-ICPermeation (**E-H**). Highest MCC was used as the overall criterion for identifying optimal parameters, with AUROC serving as a secondary criterion in cases where MCC values were similar following hyperparameter adjustment. (**A**) The number of LSTM layers was set to 1. (**B**) The hidden size of LSTM was set to 256. (**C**) The dropout was set to 0.3. (**D**) Learning rate was set to 5 × 10^−5^. (**E**) A margin of 5 was chosen. (**F**) Lambda was set to 1.0. (**G**) Learning rate of TCL was set to 0.01. (**H**) Learning rate of classifier was set to 1 × 10^−4^.

### Interpretability analysis of BLAPE-ICIdentification

To investigate how BLAPE-ICIdentification distinguishes ion channels from non-channel proteins, we applied IG (40), which estimates the contribution of each residue to the model’s output by integrating gradients from an all-zero baseline input to the actual embeddings. Due to Bi-LSTM gradient patterns, terminal residues showed higher scores than central ones in our analysis. To correct for this positional bias, we generated *N* random sequences of the same length as the target sequence to compute a positional background contribution by averaging the absolute IG scores. The residues with the top *k* normalized scores (raw divided by background) were considered potentially critical for ion channel function, including ion permeation residues, voltage-sensing regions, and other functional motifs (Figure 5A). As an example, we analyzed human gamma-aminobutyric acid receptor subunit beta-2 (GABRB2, UniProt ID: P47870), one of the five subunits of GABA-gated chloride channels that mediate the effects of GABA, the primary inhibitory neurotransmitter in the brain (54). The identified key residues (*N* = 5, *k* = 15%) were visualized in PyMOL (Figure 5B), with residues overlapping ion permeation sites highlighted in pink. This analysis yielded an MCC of 0.375.

**Figure 5.**
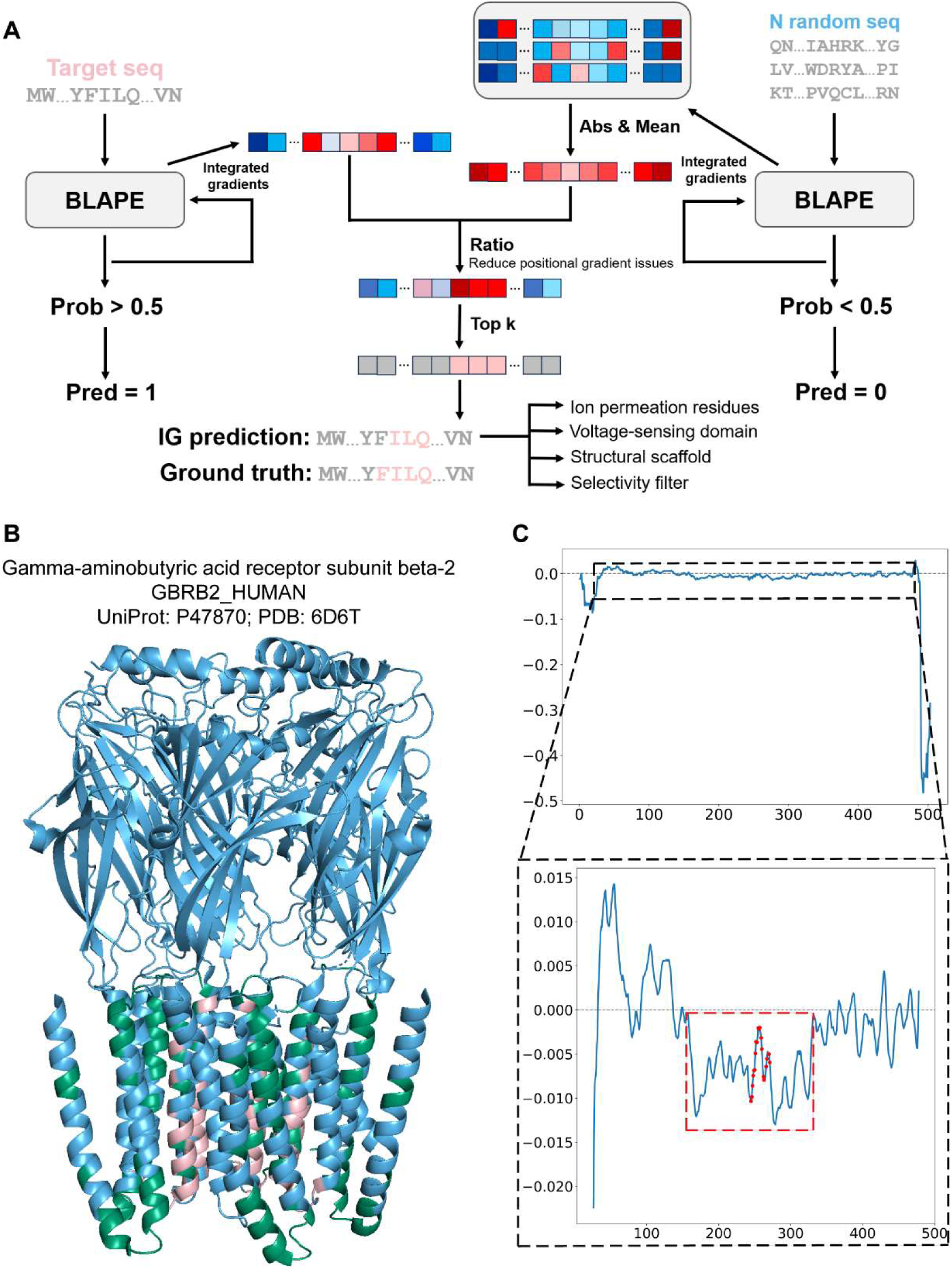
Interpretability analysis of BLAPE-ICIdentification. (**A**) Workflow of IG-based interpretability. For a sequence predicted as an ion channel, IG was applied to estimate residue contributions. Positive (red) and negative (blue) scores indicate residues promoting or opposing the prediction, respectively, with deeper colors representing stronger effects. To remove position-dependent bias introduced by Bi-LSTM, IG scores of *N* random sequences of equal length were averaged as a positional background, and normalized scores were used to identify top *k* residues potentially related to ion channel function, including ion permeation residues, voltage-sensing regions, and structural motifs. (**B**) PyMOL visualization of GABRB2 (UniProt ID: P47870). Residues overlapping known ion permeation sites are shown in pink, and non-overlapping predicted residues in green. The analysis achieved an MCC of 0.375. (**C**) Validation with amino acid deletion. Sliding-window removal of 10 residues revealed two terminal cliff regions (first and last 25 residues, outside black box) and a central drop-off (red dashed box, residues 150–330). Red dots mark known ion permeation residues.

To validate the IG results, we employed an amino acid deletion method. In detail, sliding windows of 10 consecutive residues were removed sequentially, embeddings recomputed using ESM-2, and changes in BLAPE-ICIdentification predicted probabilities were assessed. Two pronounced drop-off regions were observed at both termini, each approximately 25 residues long, consistent with IG results. After excluding these terminal “cliff” regions, we observed a distinct decrease in predicted probabilities between residues 150 and 330, suggesting this region likely harbors functional domains critical for ion channel activity (Figure 5C). It should be noted that the interpretability analysis only identifies approximate regions containing important residues, while precise identification of ion permeation residues still requires our CLAPE models.

### Amino acid composition analysis on the ICPermeation dataset

To analyze the distribution of different amino acids and their properties among ion permeation residues, we conducted an amino acid composition analysis on the ICPermeation dataset. Examining the frequencies of all 20 standard amino acids at permeation residues and non- permeation residues in the ICPermeation training set (Figure 6A), excluding unknown amino acids “X”, showed a relatively uniform amino acid distribution without obvious bias. However, four amino acids (T, G, I, and L) showed the highest frequency at permeation residues, whereas P, R, E, and K showed the highest frequency at non-permeation residues. Further scrutiny of their respective hydrophobicity of T, G, I, and L indicated that T exhibited the highest polarity, with a hydropathy index of -0.7, which was still less polar than P, the least polar amino acid among P, R, E, and K, with a hydropathy index of -1.6 (55). We thus hypothesized that weak polarity of these amino acids might facilitate ion permeation. Additionally, G had the smallest side chain, allowing for greater structural flexibility without obstructing the ion channel, whereas the rigid pyrrolidine ring of P disrupted structural continuity, potentially obstructing ion flow (56,57). For further reference, the amino acid distributions in the validation and testing sets were shown in Figure 6B, C.

**Figure 6.**
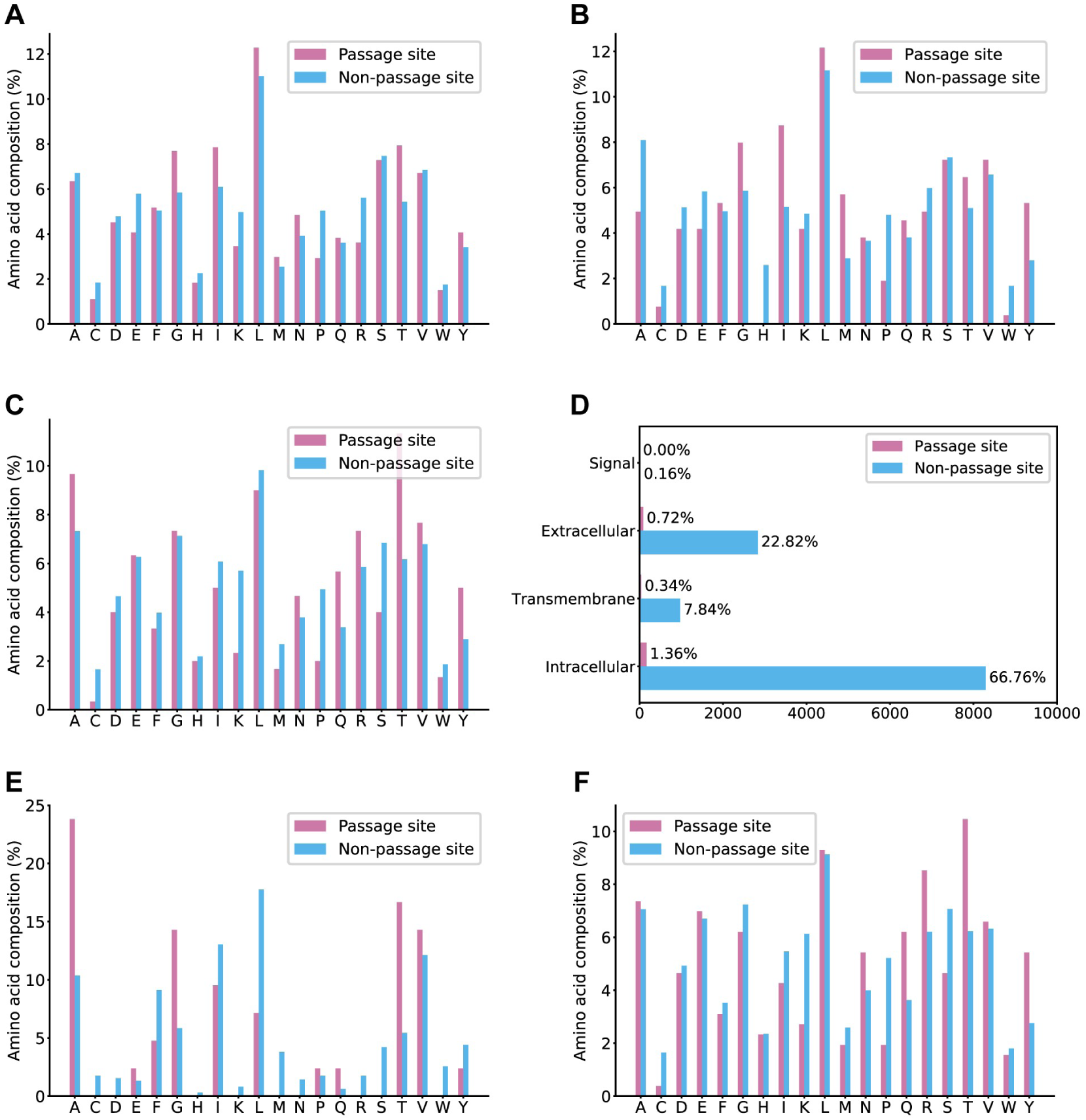
Amino acid analysis on the ICPermeation dataset. (**A, B, C**) Proportions of each amino acid at permeation and non-permeation residues in the training (**A**), validation (**B**) and testing set (**C**). (**D-F**) were analyzed on the testing set. (**D**) Distribution of permeation and non-permeation residues across different protein regions. (**E, F**) Ground truth amino acid composition at permeation and non-permeation residues in transmembrane (**E**) and non-transmembrane (**F**) regions, respectively.

To further investigate the differences in amino acid composition between transmembrane and non-transmembrane regions, we first used DeepTMHMM (41) to predict the residue-level localization of all ion channels in the testing set (Figure 6D). The results showed that only approximately 8% of the ion channel residues were located in the transmembrane region, while remaining 92% were located in non-transmembrane region. However, the proportion of ion permeation residues in the transmembrane region reached 4.2%, compared to only 2.3% in non-transmembrane regions. Next, we analyzed the distribution of amino acids in two subsets, transmembrane region subset and non-transmembrane region subset, separately (Figure 6E, F). We found that transmembrane regions showed obvious enrichment for weakly polar amino acids while show minimal presence of others (Figure 6E), whereas non-transmembrane regions showed a relatively even amino acid distribution (Figure 6F), resembling that of the entire testing set (Figure 6C). Further validation through analysis of hydrophobicity and charge distributions among ion permeation residues across the testing set, transmembrane subset, and non-transmembrane subset (Supplementary Figure 3) verified that the transmembrane region indeed exhibited obviously different physicochemical properties than non-transmembrane regions. Specifically, < 10% of transmembrane region ion permeation residues were polar, < 5% were negatively charged, and no positively charged residues were present. The remaining residues were entirely neutral.

### Comparison of CLAPE-ICPermeation and CLAPE-CaICPermeation with CaBind_MCNN and AlphaFold3 using a case study

To demonstrate the application of CLAPE-ICPermeation and CLAPE-CaICPermeation for predicting ion permeation residues, we conducted four case studies, including two ion channels from the ICPermeation testing set and two ion channels from the CaICPermeation testing set. Using PyMOL, we mapped the labels predicted by each model onto the 3D protein structures and included the results from CaBind_MCNN (23) for comparison (Figure 7). The transmembrane topology was predicted using DeepTMHMM (41), allowing us to distinguish transmembrane regions.

**Figure 7.**
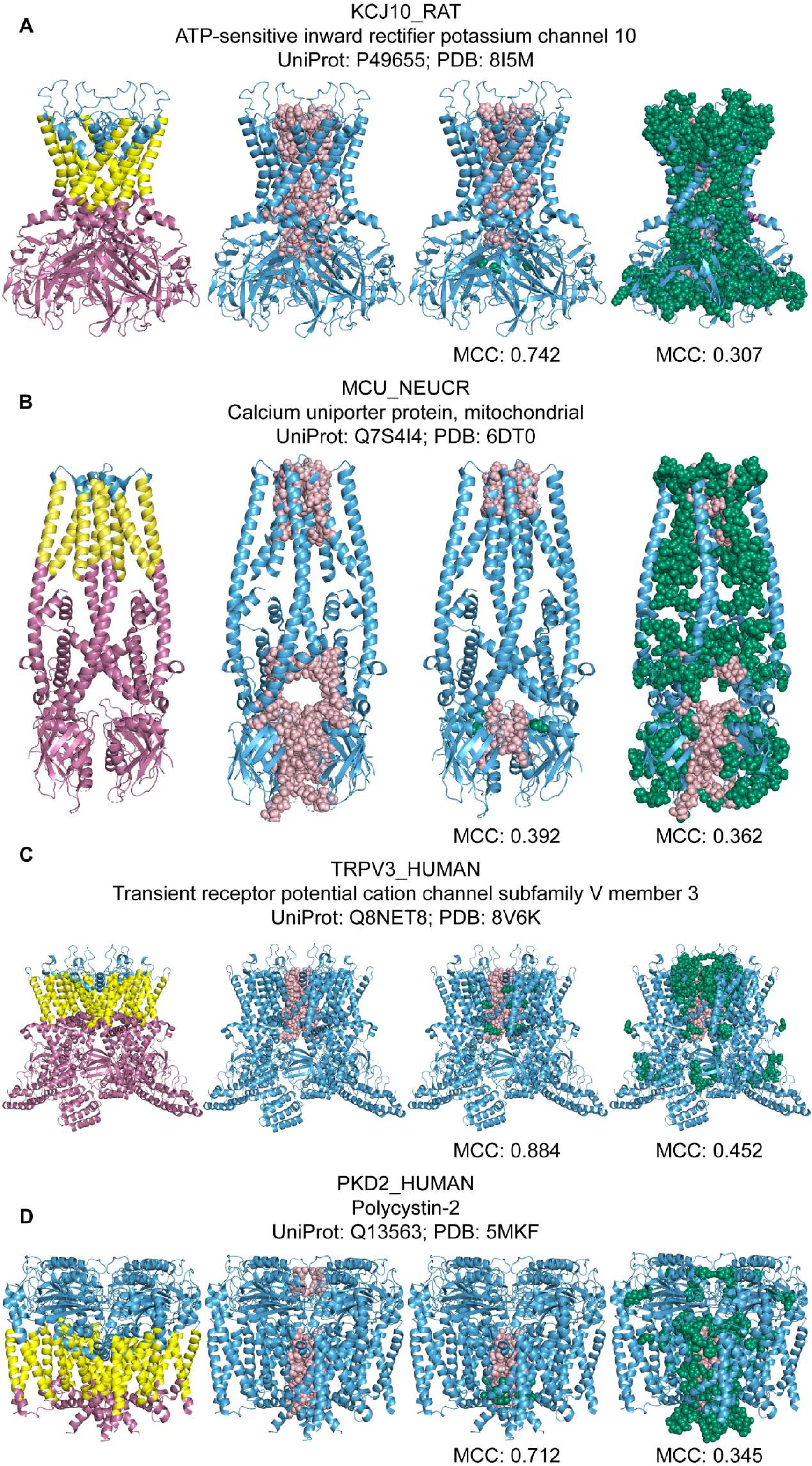
Case study of ion permeation residue prediction for P49655, Q7S4I4, Q8NET8, and Q13563. In each subfigure, the leftmost panel shows structural topology, with blue representing extracellular regions, yellow indicating transmembrane regions, and maroon indicating intracellular regions. Second panel from left displays the ground truth ion permeation residues. Third panel from left presents ion permeation residues predicted by CLAPE-ICPermeation (**A, B**) or CLAPE-CaICPermeation (**C, D**). Right panel shows ion permeation residues predicted by CaBind_MCNN. Pink, correct positive predictions; green, false positive predictions.

The first case study (Figure 7A) examined the ATP-sensitive inward rectifier potassium channel 10 of rats (KCJ10_RAT; UniProt ID: P49655; PDB ID: 8I5M), which may be responsible for the potassium buffering activity of glial cells in the brain (58). CLAPE-ICPermeation could successfully identify 14 out of 23 ion permeation residues, with few false positives (1 of 15), achieving an MCC of 0.742. In contrast, CaBind_MCNN misclassified 119 of 141 peripheral residues as ion permeation residues, resulting in a lower MCC (0.307). This characteristic was consistent with the characteristically high recall but low precision of CaBind_MCNN. In the second case study (Figure 7B), we examined the mitochondrial calcium uniporter inner membrane protein from *Neurospora crassa* (MCU_NEUCR; UniProt ID: Q7S4I4; PDB ID: 6DTO), which facilitates selective, unidirectional import of calcium ions into the mitochondrial matrix and plays a crucial role in mitochondrial calcium homeostasis (59). We found that both models showed relatively poor prediction accuracy, indicate that both models performed poorly on this case, with CLAPE-ICPermeation missing several true positives in the intracellular region (MCC = 0.392), and CaBind_MCNN generating many false positives (MCC = 0.362; Figure 7B), possibly due to the rarity of such highly selective and unidirectional transporters in the training set.

To examine our model’s accuracy in specifically analyzing calcium channels, we next conducted a case study of human transient receptor potential cation channel subfamily V member 3 (TRPV3_HUMAN; UniProt ID: Q8NET8; PDB ID: 8V6K), which is activated by warm temperatures and reportedly negatively regulates hair growth by suppressing keratinocyte proliferation and inducing premature hair follicle regression (60,61). In this protein, CLAPE-CaICPermeation correctly identified all ion permeation residues, yielding an MCC of 0.884, while CaBind_MCNN had a markedly lower MCC of 0.452 (Figure 7C). Analysis of a fourth protein (Figure 7D), polycystin-2 (PKD2_HUMAN; UniProt ID: Q13563; PDB ID: 5MKF), a nonselective cation channel involved in mechanosensation, calcium signaling, and regulation of cilium length and fluid flow that functions through homotetrameric assembly, heteromeric interaction with PKD1, or by regulating other ion channels (62,63), showed that CLAPE-CaICPermeation achieved an MCC of 0.712, primarily due to missed identification of extracellular domain ion permeation residues. In contrast, CaBind_MCNN had an MCC of 0.345, as it could recognize potential ion permeation residues in the extracellular domain, but incorrectly classified residues farther from the channel as permeation residues. These four case studies demonstrated that CLAPE outperformed CaBind_MCNN in predicting ion permeation residues.

As AlphaFold3 (43) can predict protein structures using only sequence information and specified oligomerization states, we also investigated whether ion channels could be accurately modeled in this way. For this experiment, we utilized the sequence-based protein oligomerization state prediction model, Seq2Symm (42), to infer the oligomeric states of ion channels. We then applied HOLE program (31) to identify putative ion permeation residues in the predicted structures using the same annotation criteria, and combined the predicted ion permeation residues with those obtained by AlphaFold3. We evaluated this analytical pipeline with all four of the above case study proteins (Supplementary Figure 4). Although all four proteins are tetramers, only Q8NET8 was correctly predicted by Seq2Symm as having C4 symmetry, which enabled AlphaFold3 to generate a relatively accurate structure, resulting in a relatively high MCC of 0.432. The other three proteins were predicted as having CX symmetry, indicating an uncertain oligomeric state, but likely exceeding hexameric (42).

To avoid exceeding the AlphaFold3 limit of 5000 tokens, we uniformly assigned a heptameric state for all four proteins during structure prediction. However, due to its sequence length of 968 residues, Q13563 was limited to a pentameric input. Under these incorrect oligomeric assumptions, AlphaFold3 yielded obviously compromised predictions, with MCCs of 0.006 and 0.149 for Q7S4I4 and Q13563, respectively. Interestingly, although heptameric input was also provided for P49655, AlphaFold3 could identify its correct state as tetrameric, and assembled a tetramer with the three extra copies forming an incomplete trimer. As this artifactual trimer could be manually removed during HOLE analysis, the resulting structure achieved a high MCC of 0.937. These findings indicate that providing the correct oligomeric state can enhance AlphaFold3 structural predictions, thereby improving the accuracy of ion permeation residue identification even for proteins lacking experimental structures.

### High-throughput prediction and visualization of ion channels from UniRef50 using BLAPE and CLAPE

To screen for potential ion channels in UniRef50 (36) and identify their ion permeation residues using the BLAPE and CLAPE models, respectively, we first applied the BLAPE-ICIdentification model to scan approximately 69 million UniRef50 sequences with lengths ≤3000 AA (Figure 8A). The model predicted over 217,000 known or putative ion channels. Accordingly, we used the best-performing version of CLAPE-ICPermeation to predict likely ion permeation residues in these putative ion channels identified in UniRef50. Next, we compiled a reference list of 20 representative model species, including *Homo sapiens* (human), *Mus musculus* (mouse), *Escherichia coli* (*E*. *coli*), and human immunodeficiency virus-1 (HIV-1) (see full list in Supplementary Table 6). For each protein sequence from these species predicted as an ion channel, we queried UniProtKB for PDB entries matching the corresponding UniRef ID to retrieve available 3D structures. If multiple PDBs were available, we prioritized structures determined by EM with higher resolution and greater sequence coverage. For sequences without PDB structures, we predicted their oligomeric states with Seq2Symm and calculated the token counts for AlphaFold3 as oligomeric state × sequence length. Sequences with token counts exceeding AlphaFold3’s 5,000-token limit were considered structurally unavailable. Ultimately, this process yielded structures for 2,832 predicted ion channels. Analysis of the length distributions of all UniRef50 sequences and predicted ion channels (Figure 8B) showed that both sets exhibited a long-tail distribution, in which only a small fraction of sequences exceeded 500 residues, likely representing full-length channels, while shorter sequences potentially corresponded to transmembrane fragments or misclassified membrane proteins. Notably, predicted ion channels were on average longer than UniRef50 proteins overall, which aligned with known properties of ion channels (Table 1), supporting the reliability of the model’s predictions.

**Figure 8.**
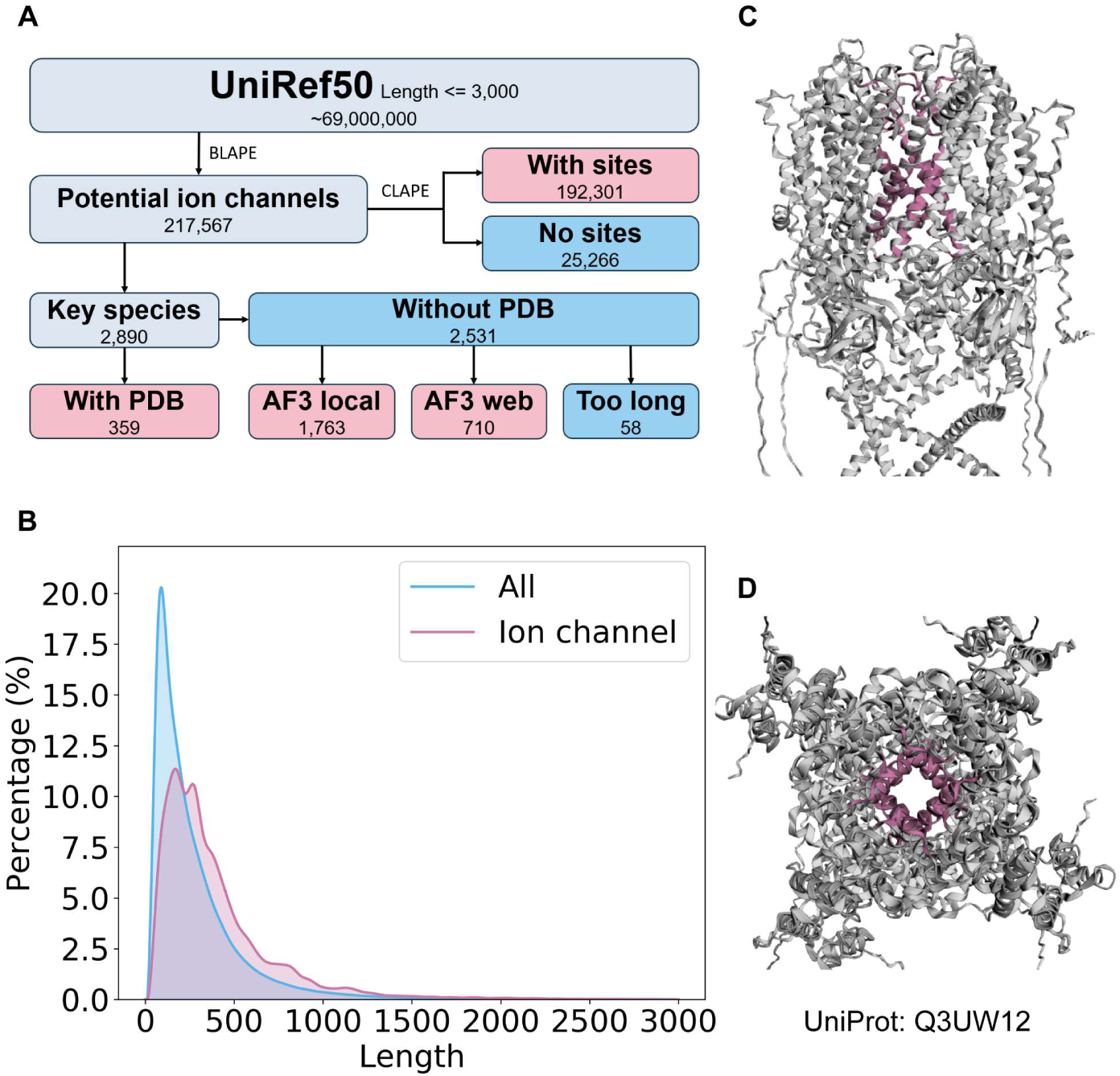
High throughput screening for potential ion channels and ion permeation residues in UniRef50. (**A**) Overview of the database screening workflow. The BLAPE-ICIdentification model first identifies 217,567 potential ion channels from among 69 million total UniRef50 sequences with length ≤3,000 AA. The CLAPE-ICPermeation model was then applied to predict ion permeation residues, yielding 192,301 sequences with predicted residues. Among 2,890 channels spanning 20 model species, 359 had experimentally determined PDB structures. AlphaFold3 was used to predict structures for the remaining 2,531 sequences, including 1,763 via local GPUs (≤2,500 tokens) and 710 via the AF3 webserver (≤5,000 tokens), while 58 remaining sequences were too long to process. (**B**) Length distributions of all UniRef50 sequences (blue) and predicted ion channels (pink) showing longer average sequence among ion channels. (**C, D**) Side and top views of a representative predicted ion channel (UniProt ID: Q3UW12), with ion permeation residues highlighted in pink.

We constructed the ICFinder interactive webserver (https://tianlab-tsinghua.cn/icfinder/) for exploring predicted ion channels and their ion permeation residues in UniRef50. The platform enables searching by UniRef IDs and filtering entries by species and confidence scores, with entries having confidence ≥0.4 defined as potential ion channels, and allows viewing amino acid sequences with highlighted predicted ion permeation residues. For ion channels with available 3D structures, we provided an additional structure viewer panel, enabling rotation and zooming. (see Supplementary Figure 5 for details). All identifiers are prefixed with “UniRef50”, highlighting the framework’s portability and scalability, which allows processing millions of sequences in new datasets within a day.

## Discussion

The present study aims to address two long-standing challenges in ion channel research. First, the continuous addition of numerous, unreviewed and unstructured protein sequences from diverse sources (e.g., environmental metagenomics and single-cell sequencing (64)) presents computational and statistical barriers for reliable, high accuracy screening of novel ion channels. Second, determining the precise path of ion permeation remains problematic for predicted ion channels lacking experimental structures. Correctly predicting the relevant ion permeation residues is therefore essential for both therapeutic development as well as rational design and engineering of ion channels (e.g., enabling artificial modulation of ion flux rates). To tackle these challenges, we curated the ICIdentification dataset, which is sufficiently large and statistically robust to serve as a benchmark for future studies of ion channel identification. Moreover, our finding that training on ion permeation data does not yield accurate predictions of ion binding, and vice versa, thus demonstrate that these functions represent distinct phenomena (Supplementary Figure 6).

As the absence of dedicated ion permeation datasets has limited the ability to demonstrate such distinctions in previous work, we constructed ICPermeation to serve as a publicly available systematic framework for studying permeation residues across diverse ion channels, which can facilitate transition to standardized, reproducible approaches for ion channel research. Using these datasets, we trained the BLAPE and CLAPE models, which subsequently outperformed comparable state-of-the-art prediction tools, including DeepPLM_mCNN (17), TMP-MIBS (20), and AlphaFold3 (43). Beyond improving prediction accuracy, we also sought to reduce the high computational burden and complexity of code associated with current GitHub implementations, which commonly hinder many biology researchers. To address these issues, we incorporated pre-computed UniRef50 predictions and launched a user-friendly webserver, ICFinder, to greatly simplify the prediction process.

A common challenge in training downstream protein models is data scarcity, particularly after stringent filtering for sequence similarity, which can compromise evaluation metrics and model generalization. Examining the effects of such stringency in our current study revealed that training with redundancy could improve MCC by ∼5% (Supplementary Table 7), suggesting this strategy might be applicable to other small-sample biomedical problems, such as rare disease or nucleic acid function predictions, where training data are often similarly limited (65,66). Additionally, given the controversy surrounding potential species-specific bias in ion channel prediction, we classified the ICIdentification testing set by species and independently analyzed large-sample species (≥100 sequences) and small-sample species (≤10 sequences; Supplementary Figure 7). These tests yielded MCCs of 0.759 and 0.815, respectively, indicating that ICFinder could capture features that are conserved across species and maintain robust performance for underrepresented species or rarely studied organisms.

Despite these advances, there remain several directions for improvement. For example, we trained a calcium-specific model in addition to general models, and our model could be extended to other ions (e.g., chloride, protons) or even non-ion channels (e.g., ATP channels) in future work. Similarly, specialized models could be developed to classify gating mechanisms, predict ion permeation direction, or regress conduction rates. Ultimately, a comprehensive set of such models may facilitate establishment of standardized “channel protein ID cards”, enabling prediction of all key properties using a single input sequence. Finally, current representations are generally limited to “amino acid-level” features, and therefore integrating “protein word-level” embeddings could benefit model performance, as these approaches have already been demonstrated to surpass amino acid-based features in small-molecule binding tasks (44).

As ion channel dysfunction underlies a wide range of diseases, identifying ion channels and their permeation residues can thus enhance our basic understanding of transport mechanisms while also guiding development of targeted therapies. By leveraging ESM-2 and curated datasets, we established robust frameworks, BLAPE and CLAPE, that provide stronger performance than other current methods for ion channel identification and permeation residue prediction. ICFinder, with the accompanying open-source code generated in this study, together provide methodological advances and practical resources for the ion channel research community, thereby helping to bridge the gap between unstructured protein sequence data and functional annotation, and opening new avenues for rational ion channel design and therapeutic discovery.

## Data availability

All datasets used for model training and performance evaluation in this study, including ICIdentification, CaICIdentification, ICPermeation, and CaICPermeation datasets, are publicly available at https://github.com/JueWangTHU/ICFinder. The homoPPI dataset can be accessed at https://github.com/TianBoxue-lab/ProteinWordwise. The datasets used by TMP-MIBS, DeepPLM_mCNN, and CaBind_MCNN are available at https://github.com/QuJing785464/TMP_MIBS/tree/main/data, https://github.com/s1129108/DeepPLM_mCNN/tree/main/FASTA, https://github.com/B1607/CaBind_MCNN/tree/main/data, respectively.

## Code availability

The code for training, testing, and inference of BLAPE-ICIdentification,

BLAPE-CaICIdentification, CLAPE-ICPermeation, and CLAPE-CaICPermeation is available at https://github.com/JueWangTHU/ICFinder.

## Acknowledgements

J.W. conducted the experiments, wrote and revised the manuscript. X.Z. assisted in the collection of datasets. X.F. assisted with webserver deployment. B.X. designed the study and assisted with manuscript preparation. B.T. designed the study, wrote and revised the manuscript.

