## Supplementary Information for "ICFinder: ion channel identification and ion permeation residue prediction using protein language models"

*To whom correspondence should be addressed:

**Supplementary Table 1.** Device of efficiency of different tasks

| Task | | Device | Efficiency |
|---|---|---|---|
| ESM-2 Embedding | ≤1,024 | GPU | $10^1$ / s |
| | >1,024 | CPU | $10^{-1}$ / s |
| Training | BLAPE | GPU | $10^2$ / s |
| | CLAPE | | $10^0$ / s |
| Testing / inference | | CPU | $10^2$ / s |

**Supplementary Table 2.** Model performance with different seeds

| | Seed | Pre | Rec | MCC | AUROC |
|---|---|---|---|---|---|
| ICIdentification | 42 | 0.926 | 0.658 | 0.778 | 0.966 |
| | 2025 | 0.957 | 0.579 | 0.742 | 0.969 |
| | 2026 | 0.880 | 0.579 | 0.711 | 0.953 |
| | 2027 | 0.958 | 0.605 | 0.759 | 0.960 |
| | 3407 | 0.880 | 0.579 | 0.711 | 0.961 |
| CaICIdentification | 42 | 1.000 | 1.000 | 1.000 | 1.000 |
| | 2025 | 1.000 | 1.000 | 1.000 | 1.000 |
| | 2026 | 1.000 | 0.800 | 0.893 | 1.000 |
| | 2027 | 1.000 | 0.800 | 0.893 | 1.000 |
| | 3407 | 1.000 | 0.800 | 0.893 | 1.000 |
| ICPermeation | 42 | 0.694 | 0.287 | 0.438 | 0.848 |
| | 2025 | 0.618 | 0.357 | 0.460 | 0.837 |
| | 2026 | 0.707 | 0.233 | 0.399 | 0.844 |
| | 2027 | 0.694 | 0.340 | 0.477 | 0.857 |
| | 3407 | 0.784 | 0.290 | 0.470 | 0.825 |
| CaICPermeation | 42 | 0.759 | 0.651 | 0.698 | 0.976 |
| | 2025 | 0.724 | 0.667 | 0.689 | 0.982 |
| | 2026 | 0.774 | 0.651 | 0.704 | 0.974 |
| | 2027 | 0.750 | 0.667 | 0.702 | 0.972 |
| | 3407 | 0.750 | 0.619 | 0.676 | 0.975 |

**Supplementary Table 3.** Model performance of BLAPE-ICIdentification on the independent testing set

| Pre | Rec | MCC | AUROC |
|---|---|---|---|
| $0.803 \pm 0.020$ | $0.495 \pm 0.021$ | $0.626 \pm 0.020$ | $0.935 \pm 0.004$ |

**Supplementary Table 4.** Model performance in 9-fold cross-validation with each dataset

| | Fold | Pre | Rec | MCC | AUROC |
|---|---|---|---|---|---|
| ICIdentification | 1 | 0.926 | 0.658 | 0.778 | 0.966 |
| | 2 | 0.885 | 0.605 | 0.729 | 0.969 |
| | 3 | 0.875 | 0.553 | 0.692 | 0.961 |
| | 4 | 0.885 | 0.605 | 0.729 | 0.956 |
| | 5 | 0.846 | 0.579 | 0.697 | 0.962 |
| | 6 | 0.852 | 0.605 | 0.715 | 0.965 |
| | 7 | 0.852 | 0.605 | 0.715 | 0.969 |
| | 8 | 0.840 | 0.553 | 0.678 | 0.969 |
| | 9 | 0.889 | 0.632 | 0.746 | 0.972 |
| | **Av.** | **0.872±0.026** | **0.599±0.032** | **0.720±0.029** | **0.965±0.005** |
| CaICIdentification | 1 | 1.000 | 0.600 | 0.772 | 1.000 |
| | 2 | 1.000 | 0.800 | 0.893 | 1.000 |
| | 3 | 1.000 | 0.800 | 0.893 | 1.000 |
| | 4 | 1.000 | 0.800 | 0.893 | 1.000 |
| | 5 | 1.000 | 0.800 | 0.893 | 1.000 |
| | 6 | 1.000 | 0.800 | 0.893 | 1.000 |
| | 7 | 1.000 | 0.600 | 0.772 | 1.000 |
| | 8 | 1.000 | 0.800 | 0.893 | 1.000 |
| | 9 | 1.000 | 1.000 | 1.000 | 1.000 |
| | **Av.** | **1.000±0.000** | **0.778±0.113** | **0.878±0.066** | **1.000±0.000** |
| ICPermeation | 1 | 0.694 | 0.287 | 0.438 | 0.848 |
| | 2 | 0.624 | 0.353 | 0.460 | 0.845 |
| | 3 | 0.679 | 0.380 | 0.499 | 0.862 |
| | 4 | 0.685 | 0.210 | 0.372 | 0.841 |
| | 5 | 0.626 | 0.373 | 0.474 | 0.869 |
| | 6 | 0.713 | 0.357 | 0.496 | 0.856 |
| | 7 | 0.648 | 0.343 | 0.463 | 0.838 |
| | 8 | 0.622 | 0.307 | 0.427 | 0.811 |
| | 9 | 0.677 | 0.350 | 0.478 | 0.854 |
| | **Av.** | **0.663±0.032** | **0.329±0.050** | **0.456±0.037** | **0.847±0.016** |
| CaICPermeation | 1 | 0.759 | 0.651 | 0.698 | 0.976 |
| | 2 | 0.810 | 0.540 | 0.656 | 0.954 |
| | 3 | 0.729 | 0.683 | 0.700 | 0.989 |
| | 4 | 0.745 | 0.651 | 0.691 | 0.987 |
| | 5 | 0.784 | 0.635 | 0.700 | 0.988 |
| | 6 | 0.774 | 0.651 | 0.704 | 0.987 |
| | 7 | 0.848 | 0.619 | 0.720 | 0.969 |
| | 8 | 0.778 | 0.667 | 0.715 | 0.983 |
| | 9 | 0.727 | 0.635 | 0.674 | 0.986 |
| | **Av.** | **0.773±0.037** | **0.637±0.039** | **0.695±0.019** | **0.980±0.011** |

**Supplementary Table 5.** Model performance with different protein language models

| | | MCC | AUROC |
|---|---|---|---|
| ICIdentification | ProtBert | 0.587 | 0.958 |
| | ESM-2 | **0.778** | **0.966** |
| CaICIdentification | ProtBert | 0.772 | 0.996 |
| | ESM-2 | **1.000** | **1.000** |
| ICPermeation | ProtBert | 0.311 | 0.815 |
| | ESM-2 | **0.438** | **0.848** |
| CaICPermeation | ProtBert | 0.622 | 0.949 |
| | ESM-2 | **0.698** | **0.976** |

**Supplementary Table 6.** Details of important species in the ICFinder webserver

| Taxonomy ID | Organism | Mnemonic | Num of predicted ion channels |
|---|---|---|---|
| 287 | *Pseudomonas aeruginosa* | PSEA | 199 |
| 562 | *Escherichia coli* | ECOLI | 429 |
| 1280 | *Staphylococcus aureus* | STAA | 38 |
| 3702 | *Arabidopsis thaliana* | ARATH | 146 |
| 4932 | *Saccharomyces cerevisiae* | YEAST | 26 |
| 5833 | *Plasmodium falciparum* | PLAF | 9 |
| 6239 | *Caenorhabditis elegans* | CAEEL | 227 |
| 7227 | *Drosophila melanogaster* | DROME | 284 |
| 7955 | *Danio rerio* | DANRE | 199 |
| 9031 | *Gallus gallus* | CHICK | 69 |
| 9544 | *Macaca mulatta* | MACMU | 24 |
| 9598 | *Pan troglodytes schweinfurthii* | PANTR | 16 |
| 9606 | *Homo sapiens* | HUMAN | 545 |
| 9823 | *Sus scrofa* | PIG | 58 |
| 9913 | *Bos taurus* | BOVIN | 78 |
| 9986 | *Oryctolagus cuniculus* | RABIT | 46 |
| 10090 | *Mus musculus musculus* | MOUSE | 227 |
| 10116 | *Rattus norvegicus* | RAT | 218 |
| 10141 | *Cavia porcellus* | CAVPO | 34 |
| 11676 | Human immunodeficiency virus type 1 | HV1 | 18 |

**Supplementary Table 7.** Model performance with or without redundant training sets on the CaICPermeation dataset

| | Pre | Rec | MCC | AUROC |
|---|---|---|---|---|
| Non-redundant | 0.622 | **0.730** | 0.667 | 0.951 |
| Redundant | **0.759** | 0.651 | **0.698** | **0.976** |

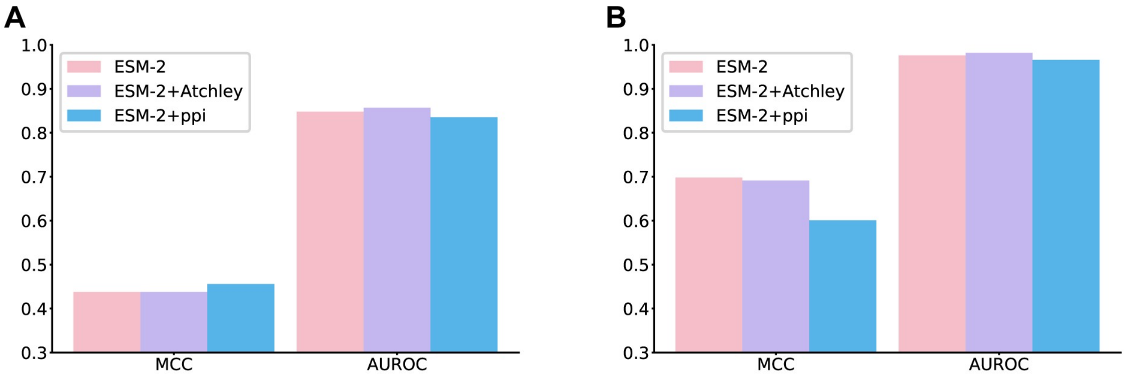


**Supplementary Figure 1.** Impact of incorporating Atchley factors and PPI information on model performance. (**A**) ICPermeation dataset, (**B**) CaICPermeation dataset.

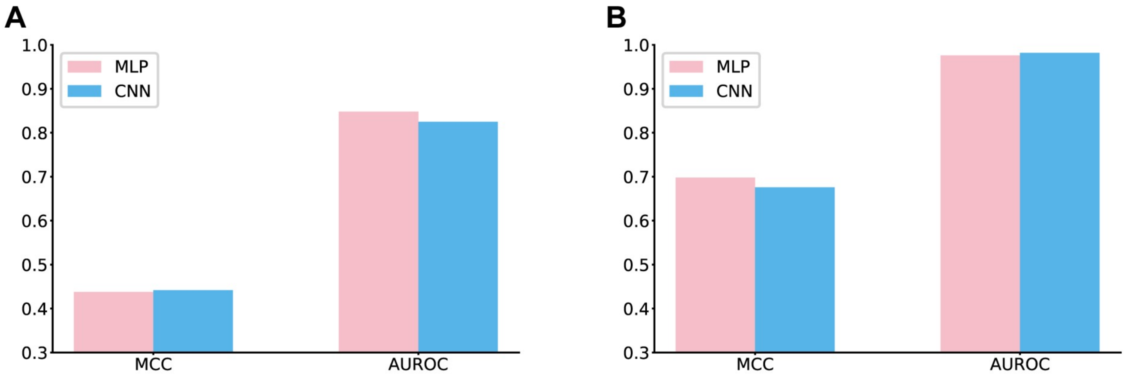


**Supplementary Figure 2.** Impact of MLP or CNN backbones on model performance. (**A**) ICPermeation dataset. (**B**) CaICPermeation dataset.

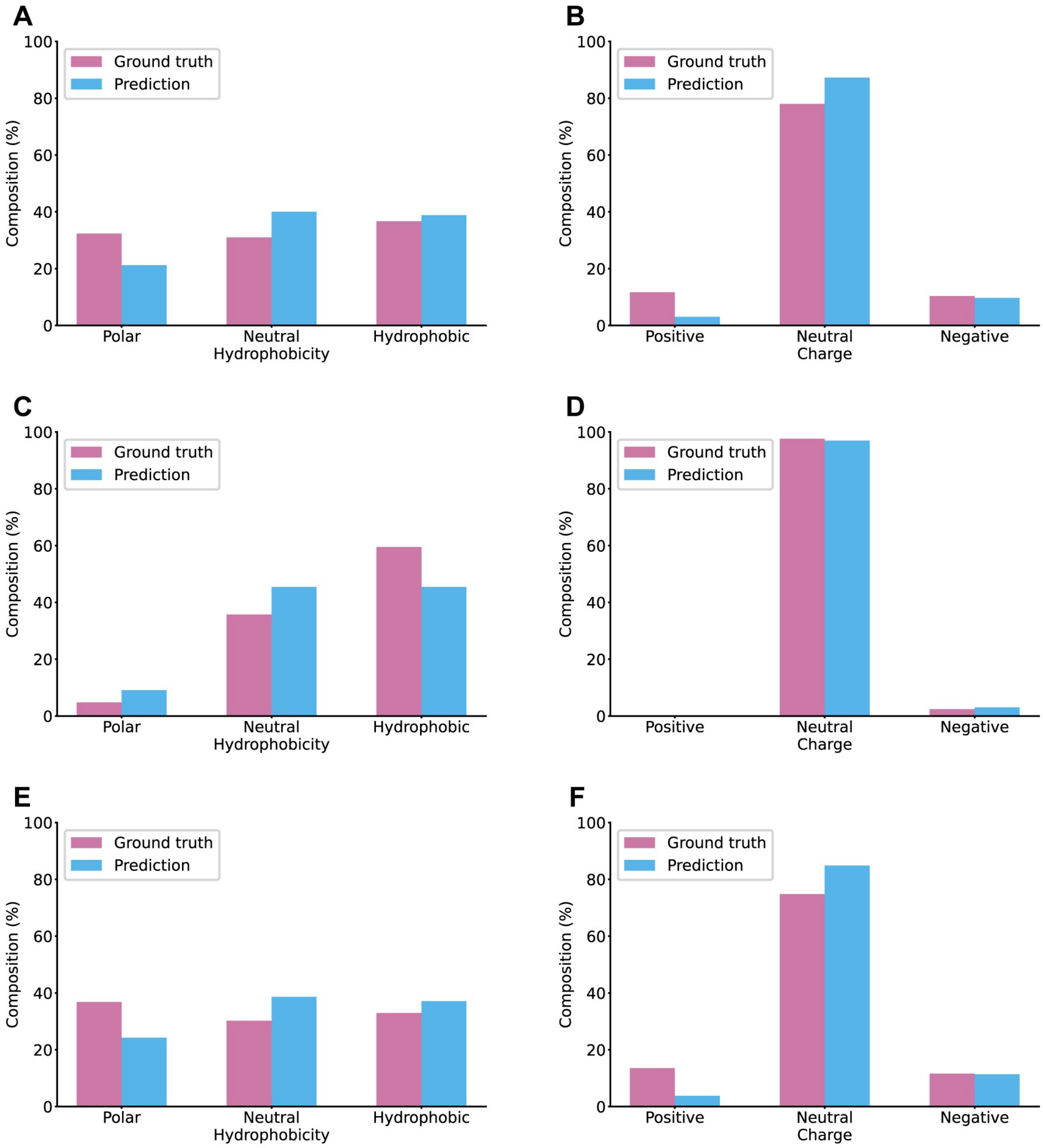


**Supplementary Figure 3.** Comparison of the true and predicted proportions of amino acid hydrophobicity (**A, C, E**) and charge status (**B, D, F**) in the ICPermeation testing set: all amino acids (**A, B**), transmembrane region (**C, D**), and non-transmembrane region (**E, F**).

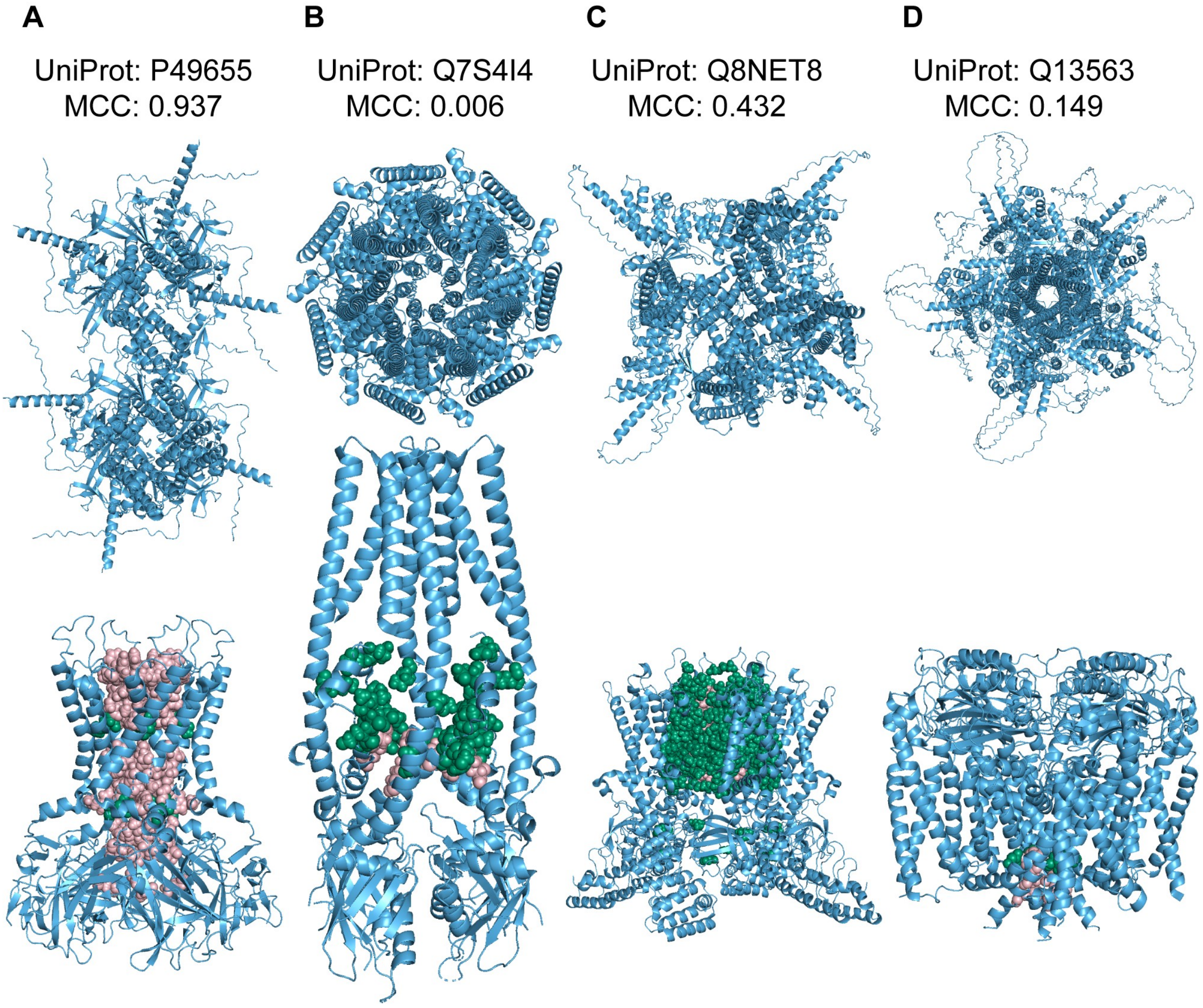


**Supplementary Figure 4.** Prediction of aggregation states and ion permeation residues for P49655, Q7S4I4, Q8NET8, and Q13563 using Seq2Symm and AlphaFold3. In each subfigure, the top panel shows the structure predicted by AlphaFold3 after providing sequence and Seq2Symm-predicted aggregation state. Bottom panel displays the predicted ion permeation residues obtained by perforating the structure with HOLE; the positions of each residue on the true PDB structure are indicated. Pink, correct positive predictions; green, false positive predictions.

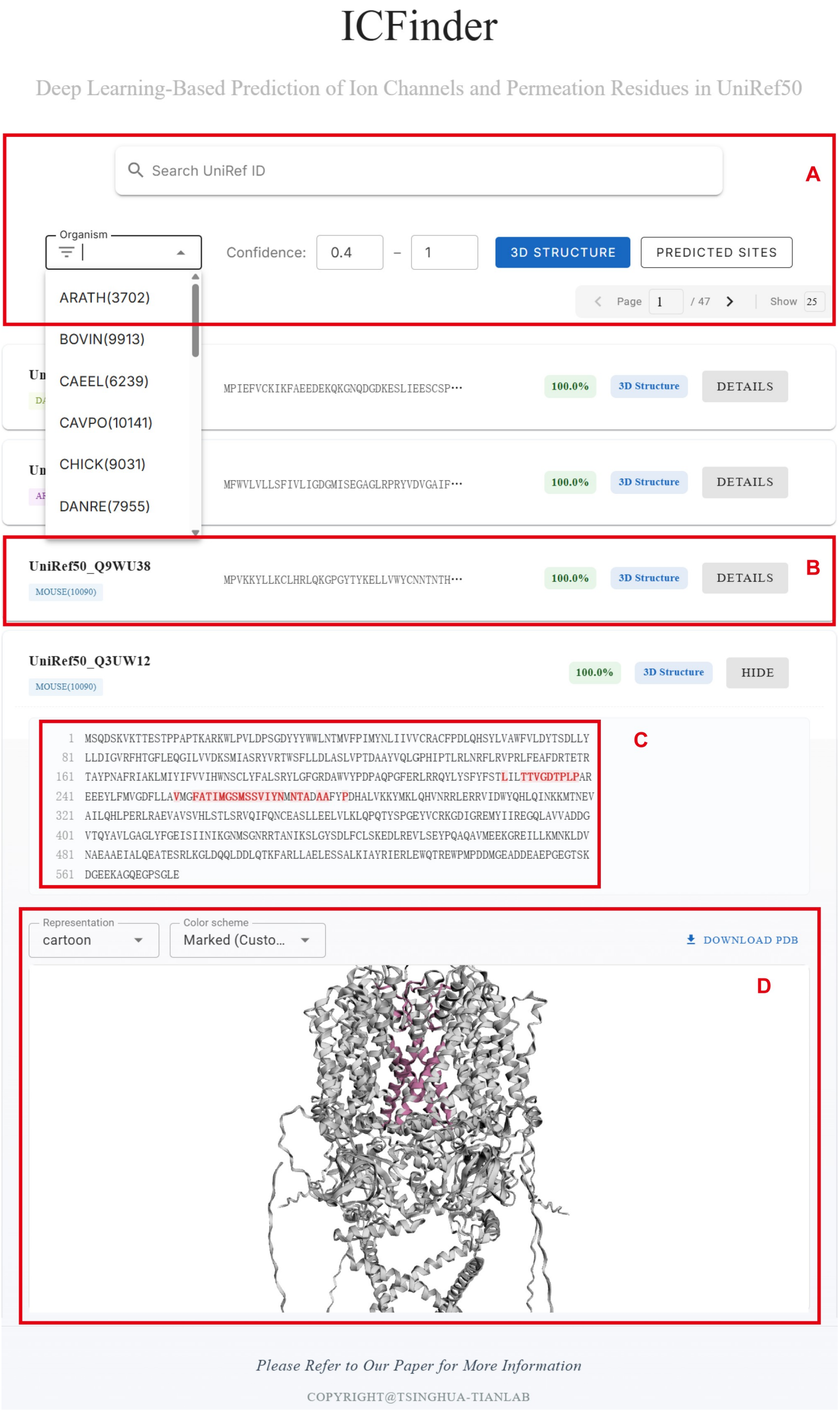

ICFinder
Deep Learning-Based Prediction of Ion Channels and Permeation Residues in UniRef50
Search UniRef ID
A
Organism
ARATH(3702)
BOVIN(9913)
CAEEL(6239)
CAVPO(10141)
CHICK(9031)
DANRE(7955)
Confidence: 0.4 – 1
3D STRUCTURE
PREDICTED SITES
Page 1 / 47
Show 25
MPIEFVCKIKFAEEDEKQKGNQDGDKESLIEESCSP…
100.0%
3D Structure
DETAILS
MFWVLVLLSFIVLIGDGMISEGAGLRPRYVDVGAIF…
100.0%
3D Structure
DETAILS
UniRef50_Q9WU38
MOUSE(10090)
MPVKKYLLKCLHRLQKGPGYTYKELLVWYCNNTNTH…
100.0%
3D Structure
DETAILS
B
UniRef50_Q3UW12
MOUSE(10090)
100.0%
3D Structure
HIDE
1 MSQDSKVKTTESTPPAPTKARKWLPVLDPSGDYYYWWLNTMVFPIMYNLIIVVCRACFPDLQHSYLVAWFVLDYTSDLLY
81 LLDIGVRFHTGFLEQGILVVDKSMIASRYVRTWSFLLDLASLVPTDAAYVQLGPHIPTLRLNRFLRVPRLFEAFDRTETR
161 TAYPNAFRIAKLMIYIFVVIHWNSCLYFALSRYLGFGRDAWVYPDPAQPGFERLRRQYLYSFYFSTLILTTVGDTPLPAR
241 EEEYLFMVGDFLLAVMGFATIMGSMSSVIYNMNTADAAFYPDHALVKKYMKLQHVNRRLERRVIDWYQHLQINKKMTNEV
321 AILQHLPERLRAEVAVSVHLSTLSRVQIFQNCEASLLEELVLKLQPQTYSPGEYVCRKGDIGREMYIIREGQLAVVADDG
401 VTQYAVLGAGLYFGEISIINIKGNMSGNRRTANIKSLGYSDLFCLSKEDLREVLSEYPQAQAVMEEKGREILLKMNKLDV
481 NAEAAEIALQEATESRLKGLDQQLDDLQTKFARLLAELESSALKIAYRIERLEWQTREWPMPDDMGEADDEAEPGEGTSK
561 DGEEKAGQEGPSGLE
C
Representation
cartoon
Color scheme
Marked (Custo…
DOWNLOAD PDB
D
Please Refer to Our Paper for More Information
COPYRIGHT@TSINGHUA-TIANLAB

**Supplementary Figure 5.** ICFinder interactive webserver for exploring predicted ion channels and their ion permeation residues in UniRef50. (**A**) Advanced search section. UniRef50 IDs can be queried using the search box, or entries can be filtered by confidence intervals to display only high-confidence ion channels or potential candidates. Users may also restrict results to sequences with available 3D structures, predicted permeation residues, or species of interest. Pagination tools are also included. (**B**) Each row corresponds to one entry, showing the UniRef ID, prediction confidence, and whether the 3D structure is available, and can be expanded using the "DETAILS" button to view detailed annotations. (**C**) Amino acid sequences are shown with predicted ion permeation residues highlighted in red. (**D**) A structure viewer is provided when available. Users can rotate and zoom the structure, adjust visualization styles and colors, or download the structure using the "DOWNLOAD PDB" button.

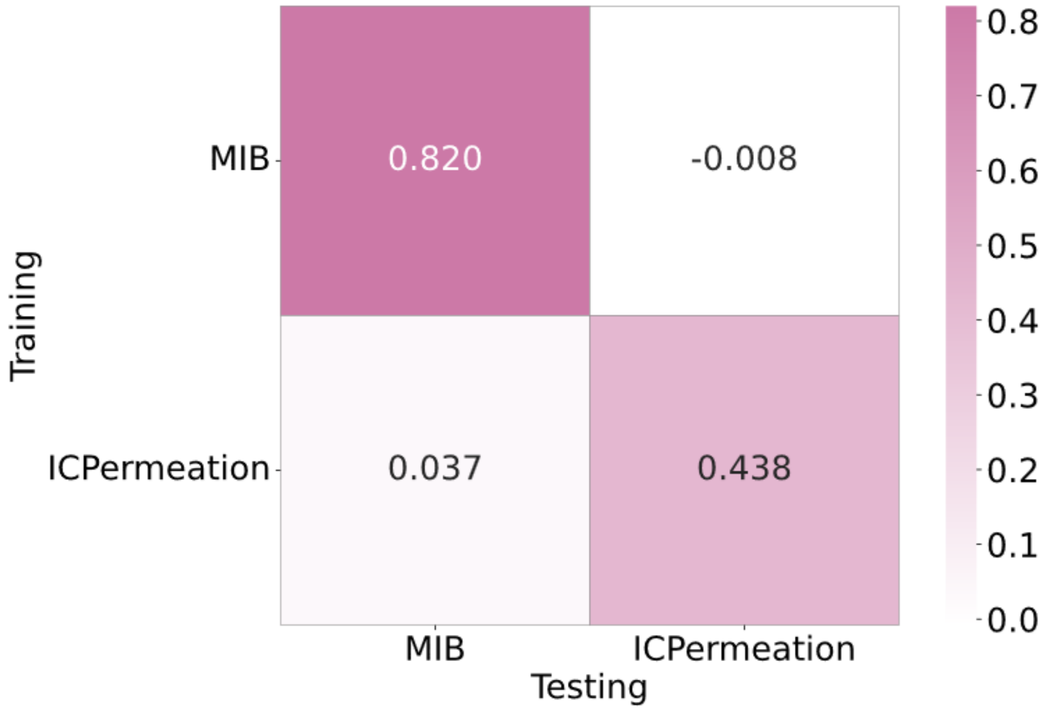


**Supplementary Figure 6.** MCC heatmap for self-training, testing, or mutual training and testing using the MIB and ICPermeation datasets.

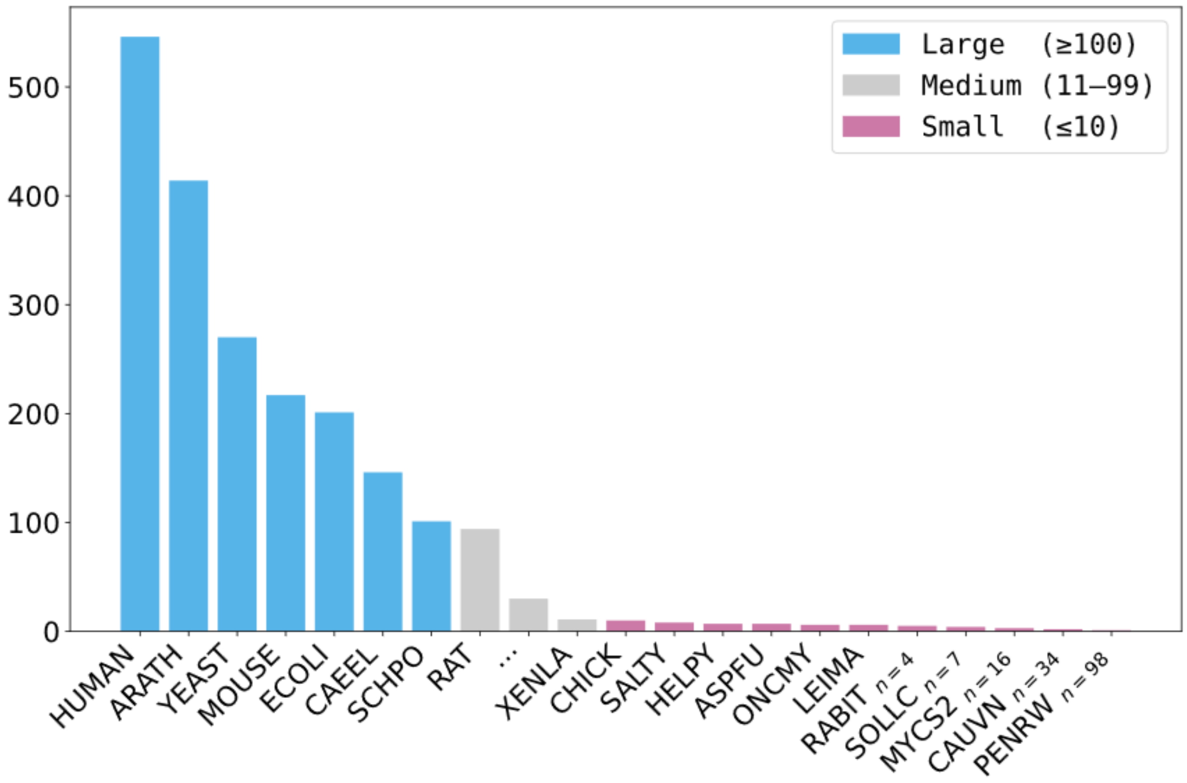


**Supplementary Figure 7.** Sequence distribution across species of the ICIdentification testing set. Large-sample species (≥100 sequences, blue), medium-sample species (11–99 sequences, gray), and small-sample species (≤10 sequences, pink). For species containing only 1–5 sequences, one representative species name is displayed with the number of such species shown in small text (e.g., RABIT $_{n=4}$).
